# On the functional role of gamma synchronization in the retinogeniculate system of the cat

**DOI:** 10.1101/2022.07.25.501474

**Authors:** Sergio Neuenschwander, Giovanne Rosso, Natalia Branco, Fabio Freitag, Edward J. Tehovnik, Kerstin E. Schmidt, Jerome Baron

## Abstract

Fast gamma oscillations, generated within the retina, and transmitted to the cortex via the lateral geniculate nucleus (LGN), are thought to carry information about stimulus size and continuity. This hypothesis relies mainly on studies carried out under anesthesia and the extent to which it holds under more naturalistic conditions remains unclear. Using multi-electrode recordings of spiking activity in the retina and the LGN of the cat, we show that visually driven gamma oscillations are absent for awake states and are highly dependent on halothane (or isoflurane). Under ketamine, responses were non-oscillatory, as in the awake condition. Response entrainment to the monitor refresh was commonly observed up to 120 Hz and was superseded by the gamma oscillatory responses induced by halothane. Given that retinal gamma oscillations are contingent upon halothane anesthesia and absent in the awake cat, such oscillations should be considered artifactual, thus playing no functional role in vision.

## Introduction

For decades, the retina has been known to generate fast gamma oscillatory activity under steady or transient photic stimulation. This phenomenon has been reported in the spiking activity of retinal ganglion cells for a wide range of species as diverse as eels (Adrian and Matthews, 1928), frogs (Arai et al., 2004; Ishikane et al., 1999, 2005), mice (Roy et al., 2017; Saleem et al., 2017; Storchi et al., 2017), rabbits (Ariel et al., 1983; Arnett and Spraker, 1981), cats (Arnett, 1975; Bishop et al., 1964; Ito et al., 2010; Laufer and Verzeano, 1967; Neuenschwander and Singer, 1996) and monkeys (Doty and Kimura, 1963). As already proposed by Adrian and Matthews (1928), such rhythmic activity is likely to emerge from cooperative interactions among cell groups (Arnett, 1975; Doty and Kimura, 1963; Laufer and Verzeano, 1967; Neuenschwander et al., 1999; Steinberg, 1966). More recently, experimental and computer-simulation studies showed that widespread gap-junctional coupling mediates synchronous activity between amacrine and retinal ganglion cells (Bloomfield and Völgyi, 2009; Roy et al., 2017; Stephens et al., 2006; Völgyi et al., 2013). These rhythmic signals are transmitted to the LGN (Arnett, 1975; Koepsell et al., 2009; Laufer and Verzeano, 1967; Neuenschwander and Singer, 1996; Storchi et al., 2017), and the cortex (Castelo-Branco et al., 1998; Saleem et al., 2017).

Retinal gamma oscillations may be essential for stimulus encoding. A robust finding supporting this hypothesis is the correlation between oscillation strength and basic attributes of light stimuli, such as size and luminance (Laufer and Verzeano, 1967; Neuenschwander et al., 1999; Saleem et al., 2017; Storchi et al., 2017). Furthermore, breaking up a continuous surface by adding a local contrast suppresses synchronization. Synchrony between oscillatory responses evoked by two contiguous bars disappears when they are separated, although the responses to the individual bars remain oscillatory (Neuenschwander and Singer, 1996). These observations in the cat have been confirmed by Roy et al. (2017) in the isolated retina of the mouse. The blockade of gap junctions or genetic ablation of connexin 36 was sufficient to suppress correlated activity among retinal ganglion cells via amacrine cells. Such manipulations also significantly impaired behavioral performance in a discrimination task (Roy et al., 2017). Furthermore, Ishikane et al. (2005) showed that, in the frog, the escape behavior triggered by a looming stimulus depends specifically on the gamma synchronization of retinal ganglion cell responses. These findings support the notion that a population-based gamma synchronization mechanism in the retina may carry spatial information that is not available in the output signal of individual ganglion cells (Koepsell et al., 2010; Stephens et al., 2006).

However, our understanding of the role of retinal gamma oscillations in visual processing is still incomplete. Most experiments have been performed with simple flashed stimuli (Laufer and Verzeano, 1967; Neuenschwander et al., 1999). It would be essential to verify whether dynamical stimuli can also be encoded by a gamma-oscillation-based mechanism. Moreover, retinal gamma oscillations in awake animals have been reported only in frogs and mice, raising the question of whether it extends to other species. Here, we address these problems by recording spiking activity and local field potentials in the retina and LGN of anesthetized and awake cats. We used dynamic stimuli, such as natural scenes and size-varying light disks. Unexpectedly, we found that gamma responses in the retinogeniculate system are strongly dependent on halothane (as well as isoflurane) and absent under ketamine anesthesia. Recordings in awake animals (N= 2 cats) revealed no signs of gamma in the LGN. Notably, in the awake cat, we often observed entrainment of the responses to the monitor refresh. These externally driven oscillations are replaced by gamma synchronized activity under halothane. Overall, our results do not support the notion that gamma responses in the retina play a role in natural vision.

## Materials and Methods

Data were obtained from 9 adult cats (2.0 to 2.5 kg) maintained within our animal facility. All experimental procedures were approved by the Ethics Committee for Animal Experimentation of the Federal University of Rio Grande do Norte (CEUA-UFRN 019/2012), Natal, Brazil.

### Experimental design

In a set of acute experiments, we recorded from the retina and the LGN of anesthetized and paralyzed cats (N= 7). In these experiments, we explored the stimulus dependencies of gamma responses under ~ 1% halothane (HALO condition). In a subset of recordings, we also investigated the effects of halothane withdrawal on gamma synchronization. Halothane was interrupted for up to 1 hour, while general anesthesia was maintained by a ketamine/ xylazine cocktail (KETA condition). In these washout protocols, halothane was eventually re-introduced (recovery transition). Since ketamine has a relatively slow clearance rate (elimination half-time: ~ 80 min; Hanna et al., 1988), it is likely that, in our experiments, ketamine still had a central action after returning to 1.0% halothane. For this reason, we refer to this recovery condition as HALO (KETA). In some control cases, designated as HALO (ONLY), we recorded from cats under halothane without immediate exposition to ketamine/xylazine. In other control cases, referred to as HALO + KETA, ketamine/xylazine was added to 1% halothane. For three recovery transitions, we used isoflurane to check if the effects of halothane extended to other halogenated, inhaled anesthetics (Hudson and Hemmings, 2019). All these experiments, which typically lasted about 72 hours (3 consecutive days), were terminal.

In a second set of experiments, two cats were trained to sit quietly for 2 to 3 hours in front of a monitor screen while their head was immobilized. This approach allowed us to obtain stable recordings from the LGN of awake cats (AWAKE condition). Subsequently, while keeping the recording electrodes in place, the cats were anesthetized, first with ketamine, then with halothane, conditions referred to as KETA (ONLY) and HALO (KETA), respectively. Thus, direct comparisons between awake and anesthesia states could be made for the same recording site.

### Surgical procedures for acute recording experiments

The cats were prepared for acute experiments by first medicating them with atropine sulfate (0.02 to 0.04 mg / kg, administered intramuscularly, i.m.) and antibiotics (ceftriaxone, 100 mg / kg, i.m.). Anesthesia was then induced by an injection of ketamine (10 mg / kg, i.m.) combined with xylazine (2 mg / kg, i.m.). After tracheotomy, the animals were artificially ventilated and maintained under halothane (1.0 to 1.2%) in a N_2_O and O_2_ mixture. In one experiment, we used isoflurane instead of halothane. The volume of the respiration pump and the respiration frequency were regularly set to keep the ventilation pressure (~ 7 mBar) and expiratory CO_2_ (~ 3.3%) within the physiological range. Body temperature was maintained at 38 °C by a thermostatically controlled heating pad. Other relevant parameters for life support (electrocardiogram, expiratory CO_2_ and pulse oximetry traces, inspiratory and expiratory volatile anesthetics, N_2_O, and O_2_ concentration levels) were continuously monitored with a Dash 3000 Patient Monitor (GE Healthcare) linked to a Smart Anesthesia Multi-gas Module. The concentration levels of halothane (or isoflurane) were estimated from the expiratory readings of the monitoring unit. Fluid losses were compensated by infusion of saline, administered intravenously at 6 ml / h.

Eye movements were blocked by i.v. infusion of pancuronium (loading dose of 0.5 mg / kg; maintenance dose of 0.25 mg / kg / h). Atropine sulfate (1 %) was applied topically to induce long-lasting mydriasis and loss of accommodation reflex. The nictitating membranes were retracted with phenylephrine (5%). The corneas were protected with contact lenses of 3 mm aperture diameter. In order to have the stimulus in focus (at a distant of 57 cm from the monitor), power correction lenses were chosen based on a refractometer (Rodenstock) readings. The optic disk and area centralis of each eye were back-projected onto the monitor screen using a fundus camera (Zeiss) and served as landmarks for referencing of the position of receptive fields. Every 8 – 12 h, the contact lenses were removed and cleaned. The clarity of the refractive media of the eyes was checked with an ophthalmoscope, and retinal fiducial landmarks were remapped on the monitor screen whenever necessary.

To obtain recordings from retinal ganglion cells, we followed the approach developed initially by Cleland et al. (1971). After opening the skin laterally to the cantus, the conjunctiva around the eyeball was cut, and the sclera exposed. A ring was sutured to the sclera, and connected to the stereotaxic apparatus with an articulated arm. This system could be moved for precise positioning of the eye. The fundus camera was used to adjust the eye’s position, according to the retinal landmarks (the area centralis and the optic disc) projected onto the screen. After opening the sclera by electrocautery, the electrodes, which were previously mounted into individual guide tubes, were inserted into the eye through a single large cannula (1.2 × 10 mm). A plate orthogonal to the eye-ring served as support for the recording device, such that no pressure was exerted on the eye. The apparatus allowed for angular rotations of the cannula in the posterior chamber. With the help of the fundus camera or an ophthalmoscope, the electrodes could thus be aimed at any desired location in the retina. This approach allowed for stable single-cell recordings over several hours. Recording sessions were discontinued when eye’s optics started to degrade, which usually occurred within 2 to 3 days.

For the LGN recordings, a single recording chamber (15 mm in diameter) was surgically cemented onto the skull across the midline. The cranium was opened, the dura removed and the cortex exposed. The electrodes were positioned into the LGN at Horsley-Clarke coordinates AP ~ 6.5 and ML ~ 9.5, corresponding to the central representation of the visual field.

### Animal preparation for chronic experiments

We developed a head-fixation system based on two chambers (9 mm diameter) implanted above the LGNs of each hemisphere for the chronic recordings. During the recording sessions, only one chamber was used to access the brain. The chambers were surgically secured onto the bone by titanium screws (Synthes standard cortex screws, 2.0 mm) and acrylic cement, under general halothane anesthesia. We allowed one month recovery period before starting with the experimental sessions. Typically a recording session was scheduled one every 1-2 weeks. Daily cleaning of the chamber with saline controlled local infections. At the end of each experiment, the cats were recovered and returned to the colony.

During the recordings, the head of the cat was fixed by the two chambers. The recording device was mounted to one of the chambers via a coupling ring adapter (screwed on top of the chamber using a thread made within it). The cats were habituated to sit in front of the computer screen with their head fixed. No additional body restraint was necessary. Wet cat food was frequently delivered as positive reinforcement during the habituation and recording sessions. This approach was based on the recording methods in awake cats developed by Bouyer et al. (1983) and had a duration of 2 to 3 hours.

We did not control for eye movements. Nonetheless, in head restrained conditions, cats tend to maintain a stable gaze to salient stimuli presented onto the monitor screen. This natural orienting behavior allowed us to obtain reliable trial-based responses, similar to the study by Kayser et al. (2003) in the visual cortex.

### Electrophysiological recordings

Neuronal activity was recorded using quartz-isolated electrodes (tungsten-platinum, 80 µm fiber, Thomas Recording). The electrodes were fitted into a customized recording device (designed by SN) made of 2 to 3 hydraulic Narishige microdrives (MO-95, Narishige Scientific Instrument Lab). For the recordings from the LGN, guide tubes were placed 5 to 7 mm above the thalamus. Electrodes were slowly moved until lamina A was reached, as indicated by robust responses to the contralateral eye. For the experiments where recordings were made directly from the eye, the guide tubes were advanced into the posterior chamber until about one millimeter above the retina. We then moved individually each electrode under visual and audio-feedback control, in order to target the ganglion cell layer.

Signals were amplified (X 1000) and band-pass filtered between 300 Hz and 7 kHz (HST ⁄ 16o25 headset, modified 32-channel preamplifier, Plexon) before being digitized at 32 kHz by a 12-bit resolution acquisition board (PCI-6071E, National Instruments). Signal display, acquisition, and storage were controlled by SPASS, a custom software written in LabVIEW (National Instruments).

Unit isolation was based on principal components and k-means clustering analysis. Quality of spike sorting was verified by several indicators, including: (1) clustering analysis scores; (2) no violation of an absolute refractory period of 2 ms as verified from inter-spike interval histograms; and (3) stability of spike amplitude and width across time. To study changes in single-cell correlated responses across anesthesia or state transitions, we always ensured that a cell pair composition (spike waveform characteristics) was preserved throughout the protocol.

### Visual stimulation

Visual stimuli were presented on a 21’’ CRT monitor (CM803ET, Hitachi) placed 57 cm away from the cat, and subtending 40° X 30° of visual angle. The refresh rate was set to 100 Hz, except for the protocols used for evaluation of stimulus entrainment) at a resolution of 1024 × 768 pixels (25 pixels corresponded to ~ 1.0° of the visual field). Stimulus presentation was controlled by the ActiveStim software (stimulus onset resolution, < 1ms).

The receptive field (RF) of each unit was mapped by presenting a set of high-contrast light bars (length, 40°; width, 0.5°) moving in 16 different directions incremented by 22.5° steps. RF maps were obtained by compiling the response histograms for each direction with a 10 ms resolution, corresponding to approximately 0.2° visual angle (Fiorani et al., 2014).

As test stimuli, we used circular disks of uniform luminance flashed on an otherwise dark screen. In order to analyze the effects of size and contrast on neuronal responses, we presented disks of 6 different sizes (2°, 6°, 10°, 14°, 18°, 22°) with three different luminance levels (1.0, 0.5 and 0.25; luminance maximum value, 40 cd/m^2^). Before the experiments, the screen monitor was gamma corrected by means of a ColorCAL colorimeter (Cambridge Research Systems). Sessions comprised ten stimulus repetitions yielding a total of 180 trials.

To assess the extent to which oscillatory patterning of responses was sensitive to dynamical changes of stimulus size, we built a stimulus paradigm in which a circular disk was varied in diameter following a random walk rule. The minimum and maximum sizes were 2° and 22°, respectively. This protocol, referred to as *size-walk*, had a duration of 7000 ms.

To test further the stimulus dependency of gamma oscillations, we used four additional types of dynamic stimuli: (1) gray-scale natural scene movies; (2) binary versions of the gray-scale movies; (3) dense binary noise refreshed at a temporal frequency from 25 to 50 Hz; (4) drifting gratings with spatial frequencies and velocities ranging from 1.25 to 2.0 cycles/° and 1.0 to 1.5 °/s, respectively. Natural scene movies consisted of movies of our laboratory recorded with a digital video camera, referred thereafter as *LabPan* and *Medikament* (the same stimuli were used by Haslinger et al., 2012). The gratings were square-wave functions and had a duty cycle of 0.3. All stimuli were presented through a gaussian aperture mask. Trials had a total duration of 5 s, with 500 ms blank before and after the 4000 ms stimulus presentation. Each stimulus condition was repeated 10 to 20 times in a pseudorandom order.

### Data analysis

Our analysis relied primarily on obtaining univariate and bivariate estimates of gamma oscillatory activity from spiking responses. This analysis was performed on single- (SUA) and multi-unit (MUA) data, independently for ON- and OFF-cells. In this study, we did not attempt to characterize cells according to their X and Y functional profiles. Nonetheless, it is known that, in cats, both cell types exhibit equally strong oscillatory responses (Ito et al., 2010). Data analysis was carried out through an integrated suite of customized programs written in LabVIEW (NES, Neurosync).

Firing rates were estimated from peristimulus time histograms (PSTH) computed with a resolution of 25 ms. Units were deemed responsive and considered for further analysis if their evoked responses to light disk (size, 22°) stimuli were significantly higher than their mean spontaneous activity. For ON-cells, this was done by comparing the trial-based firing rate in a 500 ms window immediately before and after stimulus onset using the Sign-test (*p* < 0.05). For OFF-cells, a similar procedure was carried out around stimulus offset. Joint firing rates were computed as the geometric mean of the individual rates in a cell pair.

Gamma activity was initially assessed in the time domain by computing average auto- and cross-correlogram histograms with a 1 ms resolution and a time shift of ± 256 ms. As described in Perkel et al. (1967), a shift-predictor method was employed to assess the degree of stimulus-locking of the responses. To visualize ongoing changes of the oscillatory behavior, we built sliding-window plots (step, 10 ms; sampling window size, 256 ms). As a measure of oscillation strength, we computed a modified version of the *oscillation score* (Os) proposed by Mureşan et al. (2008). In a nutshell, this metric was obtained as follows: (1) a spline function was applied on single-trial, rate-normalized correlograms to eliminate high-frequency noise; (2) an auto power spectrum of the correlograms was then computed by fast Fourier transform (frequency resolution, 1.95 Hz); (4) the peak of the power spectrum within the gamma band (30 to 90 Hz) was detected and its amplitude divided by the mean average from the whole spectrum (10 to 120 Hz). For autocorrelograms, the central peak was removed before computing its power spectrum. As a measure of Os stability across trials, we used the *confidence score* proposed by Mureşan et al. (2008).

In addition, we computed coherence estimates for combinations of SUA-SUA and SUA-MUA cell pairs taken from the same electrode or across electrodes. Mathematically, coherence is defined as:

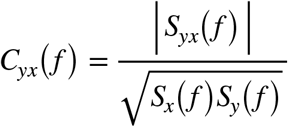

where *S*_*x*_(*f*) and *S*_*y*_(*f*)are the power spectrum estimates of the time series *x* and *y*averaged across stimulus repetitions, respectively, and *S*_*yx*_(*f*) is the cross-power of these two time series. This analysis was carried out using the multi-taper approach (Percival and Walden, 1993; Thomson, 1982) implemented in MATLAB (Mathworks) by the Chronux software package (version 2.12, available at http://chronux.org). More specifically, we used the *coherencypb* and *coherencycpb* functions after binning spike data at 1kS/s, for spike-spike and spike-field coherence analysis, respectively. Data were padded to 512 points (500 ms analysis window) before Fourier transformation. Five Slepian tapers were used. The 95% confidence intervals for the coherence estimates were determined by the jackknife method (Jarvis and Mitra, 2001). The theoretical 95% confidence limit returned by Chronux was used to assess whether coherence estimates reached a significant level. As a metric for phase-locking (PLV), we averaged the coherence phase angle obtained for individual trials using the circular statistics toolbox CircStat (Berens, 2009). The Rayleigh-test was used to evaluate whether the phase coherence had an unimodal clustering.

Besides coherence estimates, we also computed the ratio between the peak and average power from the cross-spectrum, as a quantification of joint oscillation strength for a cell pair. This metric is closely related to the oscillation score proposed by Mureşan et al. (2008) and, in our study, is referred to as OS.

### General statistics

Statistical analyses were carried out using Statview (SAS) and, unless otherwise stated, customized programs in LabVIEW and MATLAB.

We first tested the normality of each neural estimate subset (*e*.*g*., joint firing rate, Os, coherence) using the Shapiro-Wilk test for sample sizes up to 2000 or the Kolmogorov-Smirnov test for larger samples. This preliminary analysis revealed that all the data sets deviated from normality. Accordingly, subsequent hypothesis testing procedures were performed within a nonparametric framework.

For repeated measures analysis in which response estimates of the same cell pairs were compared across more than two stimulus conditions or anesthetics treatments, we used the Friedman sum rank test (e.g., Figures 5, 6, and Supplemental Figures 1 and 2). When some spectral estimates were missing due to absence of response or low firing rate (Figure 1), we used the Skillings-Mack test (Skillings and Mack, 1981), a more general form of the Friedman test that supports unbalanced data structures. When either of these two nonparametric tests produced a significant result, the Wilcoxon matched-pairs signed-rank test was used to pinpoint which pairwise groups differed significantly. The *p*-values resulting from this test were adjusted using the Bonferroni method to compensate for multiple comparisons. To assess the main and interaction effects of two factors on within cell-pair neural estimate outcomes (Figure 3), we used the aligned rank transform for nonparametric factorial analyses as developed by (Wobbrock et al., 2011) and implemented in R (https://cran.r-project.org/package=ARTool).

**Figure 1.**
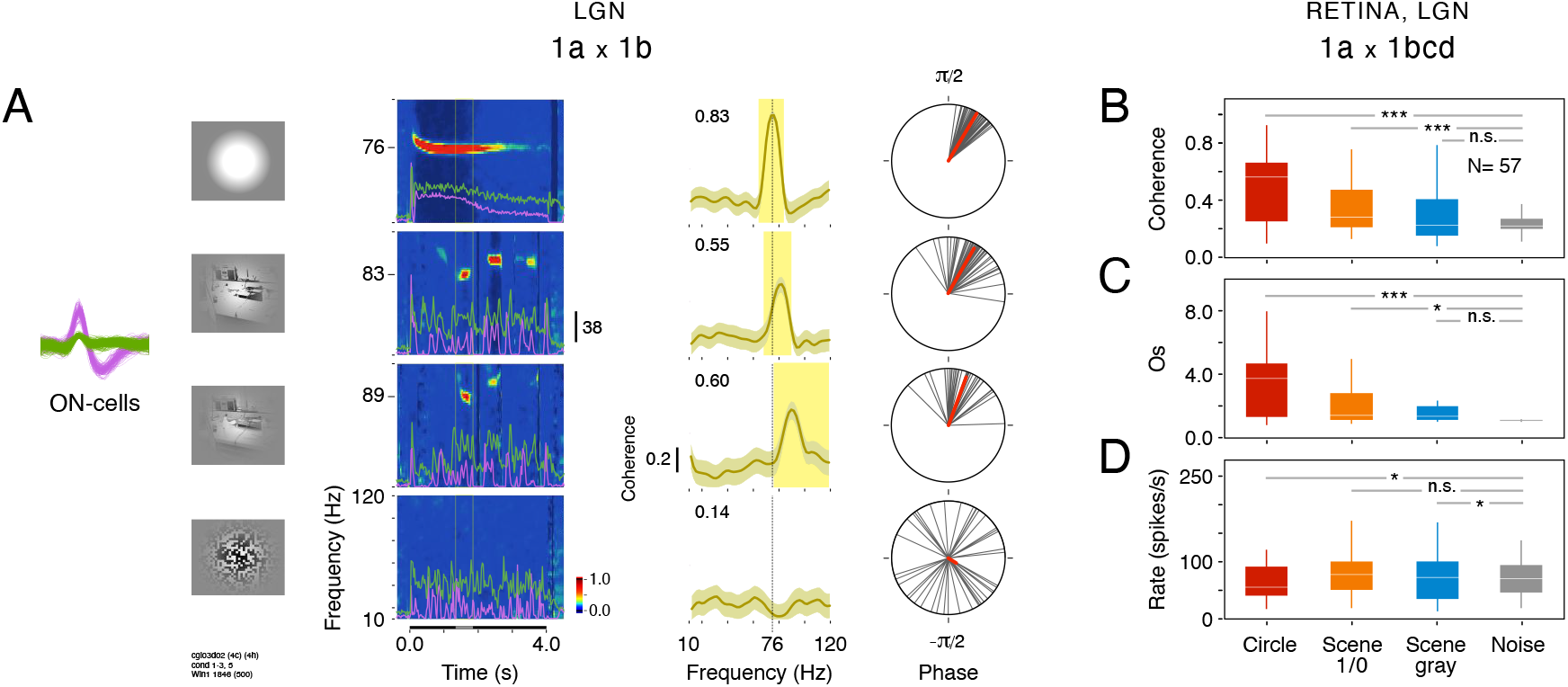
Gamma coherence is influenced by the spatiotemporal structure of visual stimuli. Single-cell and MUA recordings were made in the retina and LGN under halothane (1.0%). **(A)** Example of two ON-cells recorded from the LGN (1a x 1b, same electrode). Spike waveforms were isolated from the same electrode (*inset*). *First column*: a static light disk, binary and gray-scaled natural scene movies and dynamic noise were used as stimuli. *Second column*: sliding-window crosscorrelation time-frequency plots reveal a strong, long-lasting synchronous oscillation (peak frequency ~76 Hz, as indicated on the frequency-axis) for the large, static stimulus. In response to natural scene movies, the oscillations appeared as brief events (with peak frequencies varying between ~ 80 to 90 Hz). The noise stimulus evoked no oscillatory responses. *Third column*: average coherence functions had narrow peaks (yellow shadow, values above 95% confidence interval) consistent with the high signal/noise ratio in the correlation plots. *Fourth column*: distributions of single-trial coherence phase angles and corresponding vector average (*red line*) used as an estimation of phase-locking values. Light disk stimulus, PLV = 0.99 (mean phase angle of 60° ± 10; Z= 56.31, *p* < 0.0001, Rayleigh test); binary natural scene movie, PLV = 0.93 (61° ± 22; Z= 46.40, *p* < 0.0001); gray natural scene movie, PLV = 0.92 (70° ± 24; Z= 32.68, *p* < 0.0001); noise stimulus, PLV = 0.13 (42° ± 67; Z= 0.68, *p* = 0.51). In the time-frequency plots, stimulus duration (4s) and analysis window (500 ms, vertical lines) are represented by black and gray lines on the time-axis, respectively. *Z-scales*, relative power. Sliding-window analysis, 10 ms step and 256 ms size. PSTH scale bar, 38 spikes/s. **(B)** Comparisons of coherence distributions compiled for 57 pairs of single-cells *vs*. MUA (same electrode) in the retina and LGN according to stimulus type. Horizontal lines denote the median and the vertical lines the first and third quartiles. *** *p* < 0.0001, ** *p* < 0.001, * *p* < 0.05, n.s. non significant, Wilcoxon sign-rank test (after *post hoc* Bonferroni correction). **(C)** Distribution of oscillation score (Os) from the cross-spectrum. **(D)** Distribution of joint firing rates (Rate) estimated as the geometric mean between individual channels.

The Wilcoxon rank-sum (Mann–Whitney) and Kruskal–Wallis tests were used for two and more than two independent groups, respectively (Figure 8, 10 and 12).

Differences between population proportions were assessed by applying the likelihood ratio chi-square test (Figure 11). If the null hypothesis of equal proportions was rejected, the Marascuillo procedure (Marascuilo and McSweeney, 1967) was then used to identify the population proportions which differed significantly.

All tests were two-tailed and the level of significance was set at 0.05. Unless otherwise stated, data dispersions are shown as median and associated interquartile ranges (IQR).

## Results

### Stimulus dependencies under halothane anesthesia

To date, most studies on gamma oscillations in the retinogeniculate system have used static, uniform light stimuli. To investigate whether gamma could also encode naturalistic stimulus features, we initially decided to compare cell responses to three distinct classes of stimuli: static light flashes, natural scene movies and dynamic dense noise. For these initial experiments, anesthesia was maintained at 1.0 % halothane.

Figure 1A shows a representative example of responses from two ON-cells in the LGN. In line with a previous single-cell study (Ito et al., 2010), a large flashed light stimulus induced a strong, long-lasting synchronous oscillation for both cells (time-frequency plot). This oscillatory patterning had a single gamma component at 76 Hz and was tightly phase-locked across channels, as indicated by the high amplitude peak (0.83) in the average coherence function. In response to natural scenes movies, however, gamma oscillations appeared as brief events, lasting for a few hundred milliseconds. During these epochs, the synchronized oscillations were strong, but not as much as when induced by large static stimuli (average coherence of 0.55 and 0.60, at 83 Hz and 89 Hz, for binary and gray stimuli, respectively). It is likely that oscillations were induced only when the RFs were simultaneously exposed to continuous surface portions of the image. This occurred intermittently in our natural scene movies. Such a conjecture is reinforced by the fact that dynamic, binary dense noise triggered no oscillatory patterning, but evoked firing rates comparable to the other conditions. Note also that the trial-based coherence phase angles get progressively more dispersed, from the static stimuli to the binary noise. This overall trend was consistent within a sample of 57 pairs of ON-cells recorded from the retina and LGN of 2 cats (Figure 1B-D). A significant stimulus-dependent modulation was found for both coherence (*MS*_3_ *=* 12.68, *p* = 0.0054, Mack-Skillings test) and Os (*MS*_3_ = 24.69, *p* = 1.79 × 10^−5^). The coherence median obtained for the static light disk stimulus (0.59, IQR = 0.4) was about twice that obtained for natural scenes (binary: 0.29, IQR = 0.29; gray: 0.22, IQR = 0.22). This difference was close to the statistical significance threshold (disk stimulus *vs*. binary natural scene, *p* = 0.069; disk stimulus *vs*. gray natural scene, *p* = 0.036; Wilcoxon signed-rank test with *post hoc* Bonferroni correction). Significance was clearer, however, when similar comparisons were performed on Os values (disk stimulus *vs*. binary natural scene, *p* = 0.0008; disk stimulus *vs*. gray natural scene, *p* = 0.016). Coherence and Os median values for the dense noise stimulus were significantly lower than for any other stimulus classes (see asterisks symbols in Fig. 1. B-C). Firing rate varied across stimulus conditions (Figure 1D, *MS*_3_ *=* 14.05, *p* = 0.0028), but showed no relationship to gamma.

The results described above are compatible with the idea that gamma coherence encodes the continuity of surfaces. To test this idea more systematically, we built light spot stimuli that changed in size according to random-walk sequences, each repeated 20 times. In the representative example shown in Figure 2, the size-changing stimulus was centered at the proximity of the RFs of ON and OFF-cell groups recorded from the same electrode. With this spatial configuration, the stimulus covered intermittently the RFs, leading to activation/ inhibition of the ON/OFF cells. Brief episodes of gamma appeared in alternation for the ON and OFF-responses. The power and duration of oscillatory events followed the instantaneous size of the stimulus. Interestingly, gamma frequency was non-stationary.

**Figure 2.**
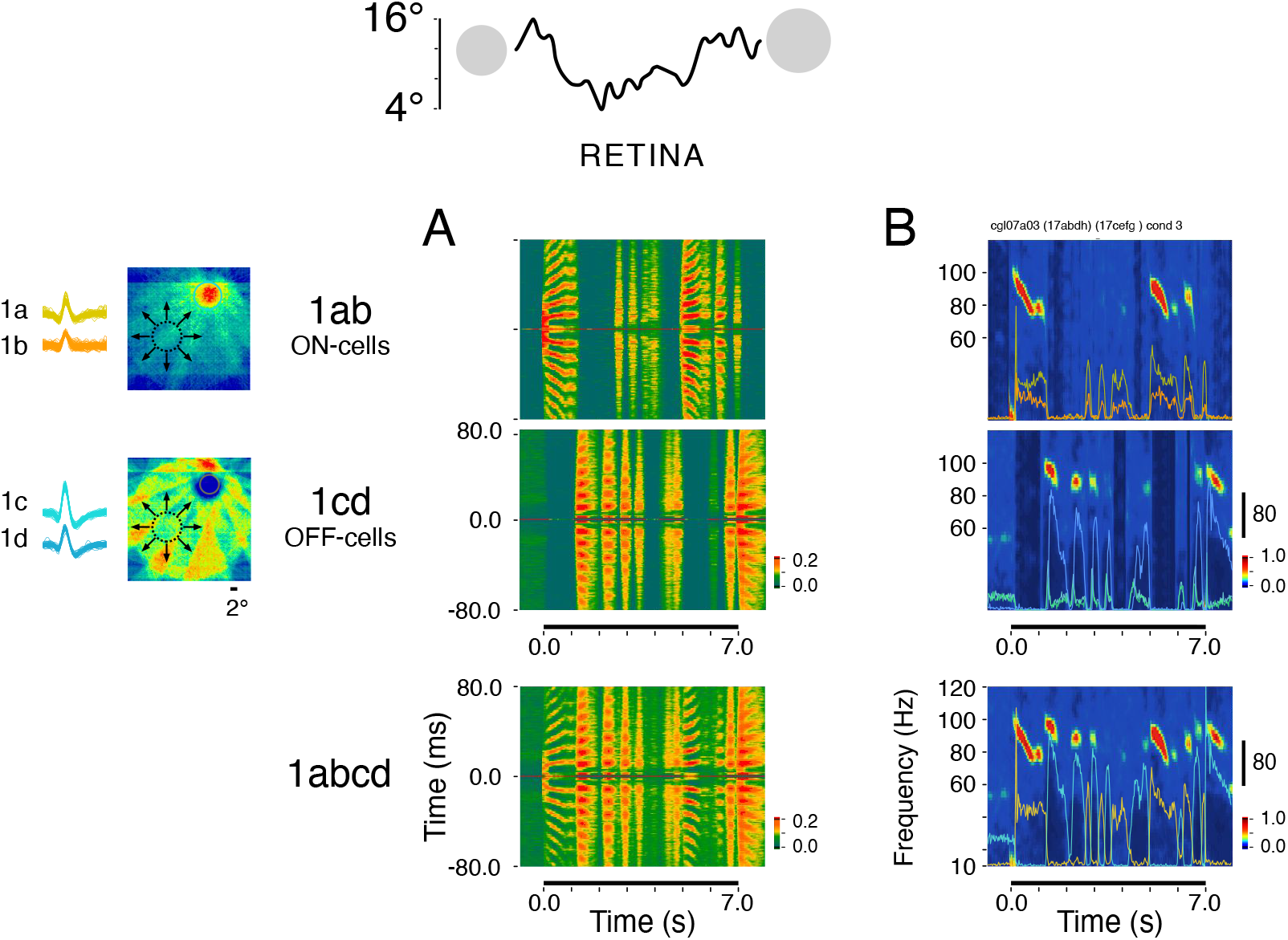
Gamma responses of both ON- and OFF-cells can follow fast stimulus dynamics. Recordings were made in the retina under halothane (1.0%). **(A)** Sliding window autocorrelograms obtained for a group of ON-cells (1ab) and a group of OFF-cells (1cd) recorded simultaneously from the same electrode. *Top inset:* stimulus consisted of a light disk continuously varying in size from 4° to 16° in random-walk manner. *Left insets:* receptive field (RF) maps for each of the two cell groups are shown together with a schematic representation of the stimulus. Notice that the stimulus was not aligned with RFs center. This configuration led to an intermittent covering of the RFs when the stimulus expanded or retracted. Response activation/suppression are mapped as warm/cold colors. RFs of the two cell groups were overlapping; as denoted by the colored circles (ON-cells, *yellow-orange*; OFF-cells, *cyan-blue*). Scale bar, 2°. Autocorrelation time shift, from −80 to 80 ms. Z-scales, autocorrelograms were normalized by firing rates. **(B)** *Upper an middle panels:* time-frequency plots of the same data shown in (A). Traces represent the PSTH of individual cells with color matching their respective spike waveforms. *Lower panel:* superposition of the time-frequency plots obtained separately for the ON- and OFF groups. Traces represent joint PSTHs (*orange*, ON-group; *blue*, OFF-group of cells).

Multiple resettings in frequency and phase are evident in the combined autocorrelation plots derived for the ON- and OFF-responses (Figure 2A). Note that gamma dynamics depended on the activation of the cells, but they were clearly not a mere effect of increase/decrease of firing rates. For example, at the beginning of the trial, the ON-responses were relatively stable, although oscillation frequency decays steadily (Figure 2B). These results reinforce the notion that gamma synchronous oscillations may provide an encoding mechanism complementary to rates. Moreover, our data show that gamma responses for the ON and OFF channels can be triggered independently.

Previous studies in different species reported that gamma in the retina is modulated by stimulus size (Ishikane et al., 1999; Neuenschwander et al., 1999) and luminance (Laufer and Verzeano, 1967; Storchi et al., 2017). Here we examined the possibility of interactions between these two parameters. The time-frequency plots of Figure 3A show examples of response profiles obtained for two ON-cells recorded from independent electrodes in the retina. RFs were overlapping, indicating that the two sites were nearby. For a large stimulus (22°), at maximum relative contrast (1.0), synchronous oscillations were strong (Os= 6.36) and persisted over the whole stimulus presentation of the stimulus. Note that, for all conditions, a pronounced decay of oscillation frequency appears right after stimulus onset, independently of the oscillation strength. This feature was always found in our data and is ubiquitous in many studies on gamma oscillations, not only in the retinogeniculate system but also in the primary visual cortex (Brunet et al., 2015; Castelo-Branco et al., 1998; Murty et al., 2018; Neuenschwander et al., 1999; Peter et al., 2019). A minimal stimulus size, larger than the center region of the receptive fields, was required to evoke a robust oscillation. At maximal luminance, we estimated the size threshold to be around 6°, consistent with previous data (Neuenschwander et al., 1999). Stimulus luminance had a marked influence on the duration of the oscillations. For low contrast stimuli, oscillatory responses were relatively brief, corresponding to the epochs of high firing rates (see accompanying PSTHs). The fact that a minimum stimulus size is needed for triggering the oscillatory responses reinforces the idea that gamma oscillations in the retina are not generated by individual retinal ganglion cells, but requires interactions across a cell population.

**Figure 3.**
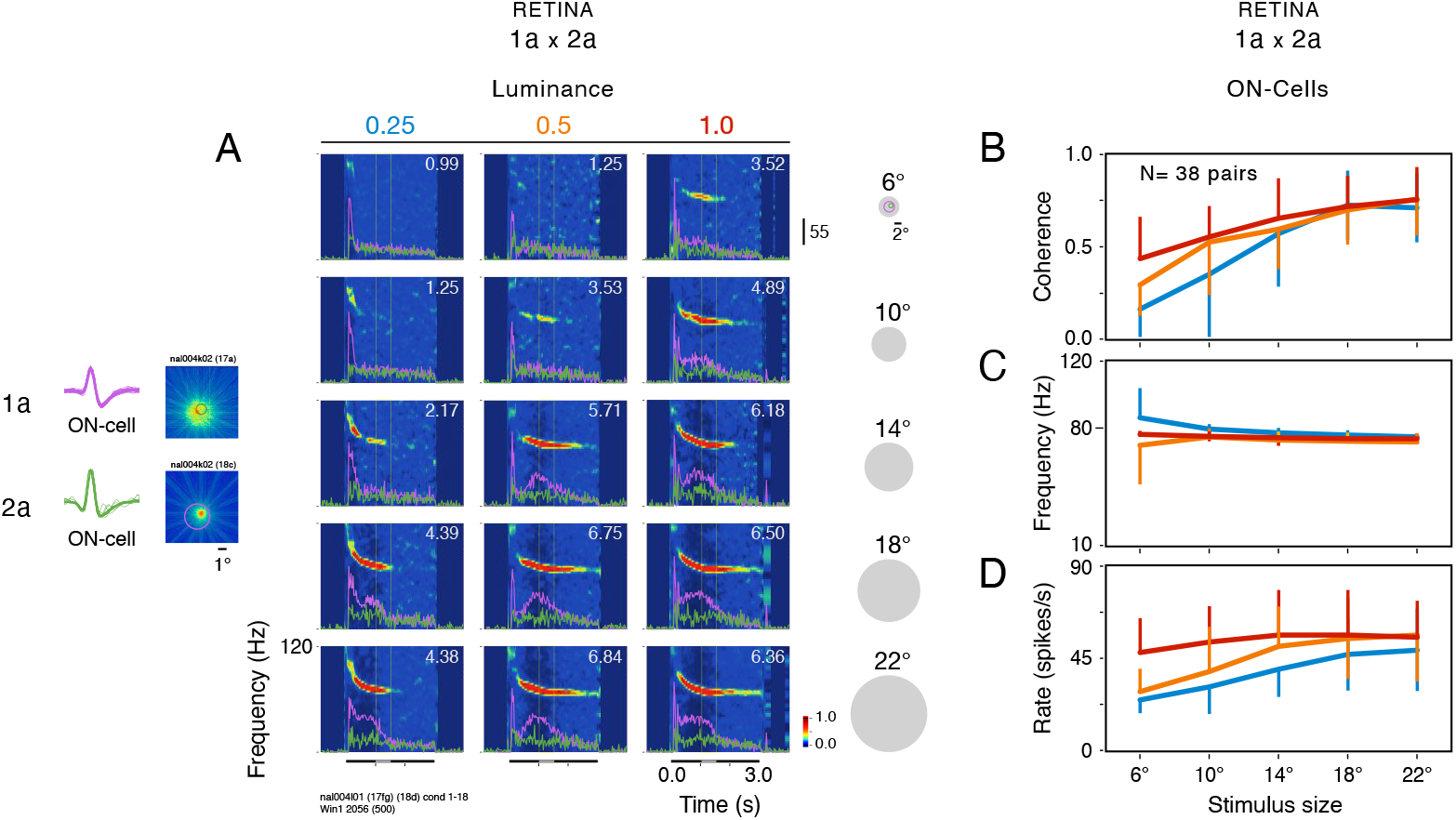
Stimulus size and luminance modulate gamma in single-cells. Recordings were made in the retina under halothane (1.0%). **(A)** Cross-correlation time-frequency plots as function stimulus and luminance. Data obtained for three different luminance levels (blue, 0.25; orange, 0.5; red, 1.0, maximum value of 40 cd/m^2^). Light disk of 6°, 10°, 14°, 18° and 22° size were centered over the RFs (scale bar, 2°). Oscillation scores (Os) are shown on the upper-right corner of the panels. **(B)** Population coherence for 38 pairs of ON-cells in the retina (1a x 2a, cross-electrodes). Error bars represent the standard deviation (SD) around the mean. **(C)** Frequency at gamma peak obtained from cross-spectrum. **(D)** Joint firing rates (Rate) estimated as the geometric mean between individual channels.

Figure 3B-D shows how coherence, frequency and firing rate are related to stimulus size and luminance. Data were obtained for 38 cross-electrode cell pairs in the retina. Nonparametric factorial analysis based on aligned rank transformations revealed a significant interaction effect of stimulus size and luminance on coherence (F_10,37_ = 9.8, p < .0001). Main effects were also highly significant (size: F_5,37_ = 374.5, *p* < 0.0001; luminance: F_2,37_ = 50.0, *p* < 0.0001). Similar results were encountered for Os (Size*luminance interaction: F_10,37_ = 6.7, p < 0.0001; Size main effect: F_5,37_ = 269.5, *p* < 0.0001; Luminance main effect: F_2,37_ = 17.0, *p* < 0.0001). Interestingly, an increase in gamma strength as a function of stimulus size was also found in monkey V1 (Gieselmann and Thiele, 2008; Jia et al., 2013; Ray and Maunsell, 2010; Zhang and Li, 2013). In the cortex, however, gamma frequency decreased as a function of stimulus size and increased as a function of contrast (Jia et al., 2013). These effects can add up to 30 Hz. In our data, we observed a small but significant change (less than 5 Hz) in frequency for both size (F_4,37_ = 36.9, *p* < 0.001) and luminance (F_2,37_ = 37.3, *p* < 0.0001). Also, the effect of size on frequency depended on the level of luminance (F_8,37_ = 8.0, *p* < 0.0001). In this set of experiments, there was a significant correlation between coherence and firing rate (Pearson *r* = 0.788, *p* < 0.0001). This relationship is likely to be explained by the fact that the stimulus dependency of rate was more pronounced at a later stage of the responses, which corresponded to the sustained epoch of the gamma oscillation.

### Effects of anesthesia

A critical issue, which is still unresolved, is how anesthesia affects gamma synchronization in the retina and its feedforward propagation via the LGN. Anesthesia could profoundly impact neuronal dynamics, leading to false inferences about the function of gamma in early visual processing. We conducted experiments to address this problem by temporarily interrupting halothane while maintaining general anesthesia through ketamine. To our surprise, we found that gamma oscillations are strongly dependent on halothane concentration levels. When halothane reached concentrations close to zero, the oscillatory patterning of the responses was abolished. This dose-dependent effect was insensitive to ketamine.

Figure 4 shows the results obtained in one of the three halothane-washout experiments for which response evolution was quantified on a minute basis. LGN MUA responses to a large light stimulus were first recorded under 1% halothane. We then administered ketamine (10 mg / kg, i.m.), and after 10 min, we switched off the halothane vaporizer. During the next 30 min, responses were continuously recorded. For analysis, data were compiled in consecutive sets of 20 trials, with incrementing steps of 10 trials (analysis window, 750 ms; inter-trial interval, 15 s). Over this period, exhaled halothane concentration decreased progressively from 1.0% to 0.1%. The power spectrum of time-resolved cross-correlograms

**Figure 4.**
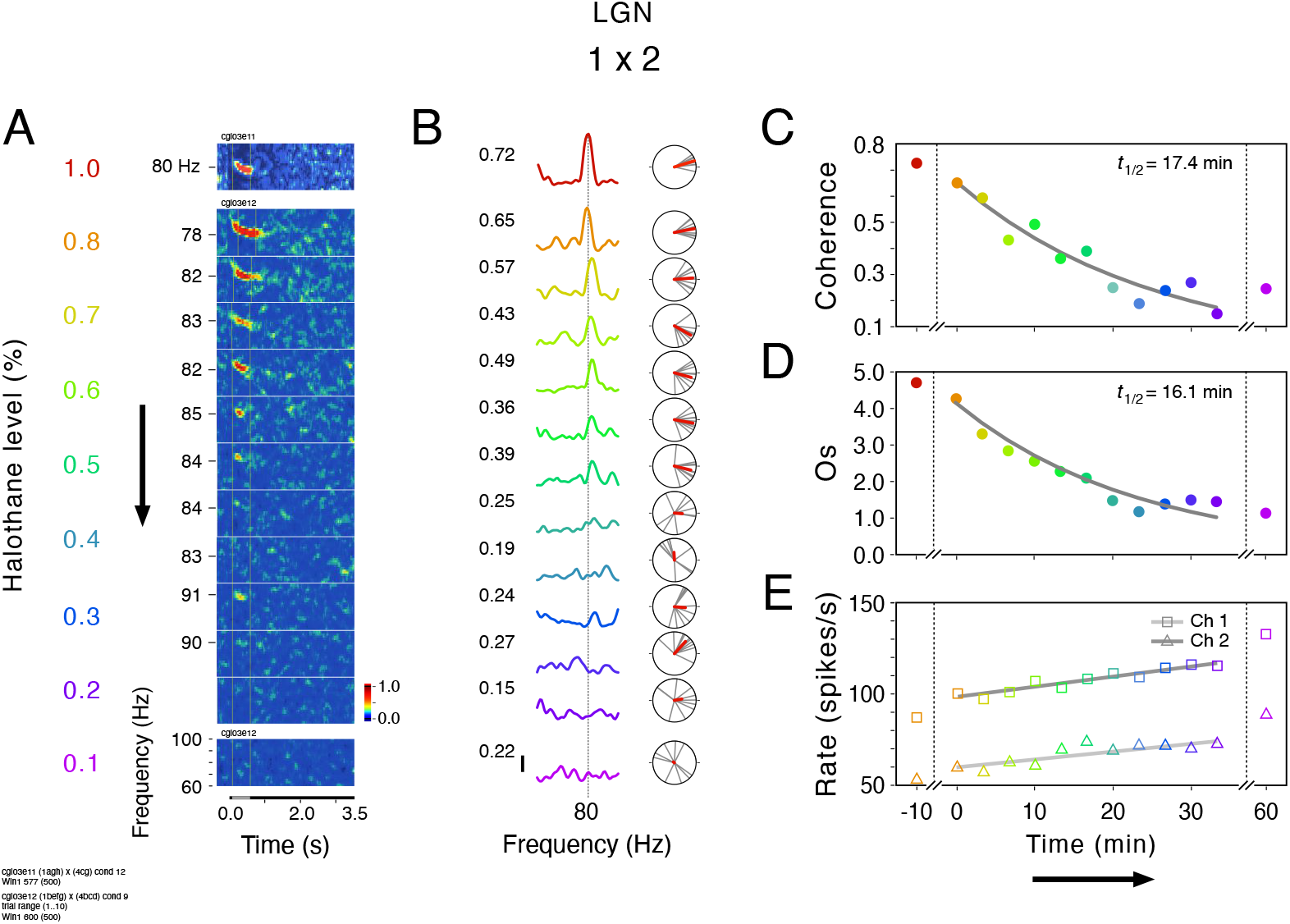
Washout of halothane drastically reduces stimulus-evoked gamma coherence. MUA recordings were made in the LGN in response to a large light disk. **(A)** Cross-correlation time-frequency plots for progressively decreasing concentration levels of halothane (from 1.0% to 0.1%, *red* to *magenta*). Data was compiled in sets of 20 trials, with incrementing steps of 10 trials (analysis window, 750 ms; inter-trial interval, 15 s). **(B)** Average coherence functions and phase distributions. Coherence peak values are indicated on the left. Scale bar, 0.2. Same color-code scheme of (A). **(C)** Decay of coherence as a function of halothane withdrawal time. A half-life of 17.4 minutes was obtained from fitting an exponential function to the data (*gray line*). **(D)** Same as in (C) but for Os metric (estimated *half-life* = 16.1 min). **(E)** Same as in (C) but for firing rates of individual channels (*gray lines*, linear fits).

(Figure 4A) and coherence functions (Figure 4B) show that stimulus-evoked gamma synchronization dropped dramatically in concomitance with the washout of the anesthetics. As plotted in Figure 4C-D, coherence and Os exhibit a shallow exponential decrease, with a lifetime of ~ 17 min. Notably, coherence did not reach a significant level below ~ 0.3% halothane (data points in blue). This dose-dependent effect could not be attributed to a decrease in responsiveness. Over the halothane washout period, firing rates showed a small monotonic increase of about 10% (Figure 4E) and were negatively correlated with coherence (channel 1, *r* = −0.85, *p* = 0.001; channel 2, *r* = −0.94, *p* < 0.0001) and Os (channel 1, *r* = −0.80, *p* = 0.0028; channel 2, *r* = −0.92, *p* < 0.0001).

Re-establishing halothane to its initial, steady-state concentration of 1.0% led to a restoration of the oscillatory responses (Figure 5). This analysis was performed for a total of 121 cell pairs (108 LGN x LGN pairs and 13 retina x retina pairs), for which SUA recordings were stable enough to follow cell responses across the entire HALO # KETA # HALO (KETA) protocol. Independent of a certain variability in coherence strength and frequency across cell pairs (Figure 5A), there was a consistent and clear-cut drop in coherence upon withdrawal of halothane (HALO # KETA transition). Figure 5B shows how the initial correlation between coherence and OS (Pearson *r* = 0.84, *p* < 0.001) changes under ketamine and reappears with a similar profile when halothane was re-established (*r* = 0.84, *p* < 0.001). The same trend occurs for the Os *vs*. rate relationship. Note, however, that the correlation between the two variables became stronger for HALO (KETA) (r = 0.66, p < 0.001) than HALO treatment.

**Figure 5.**
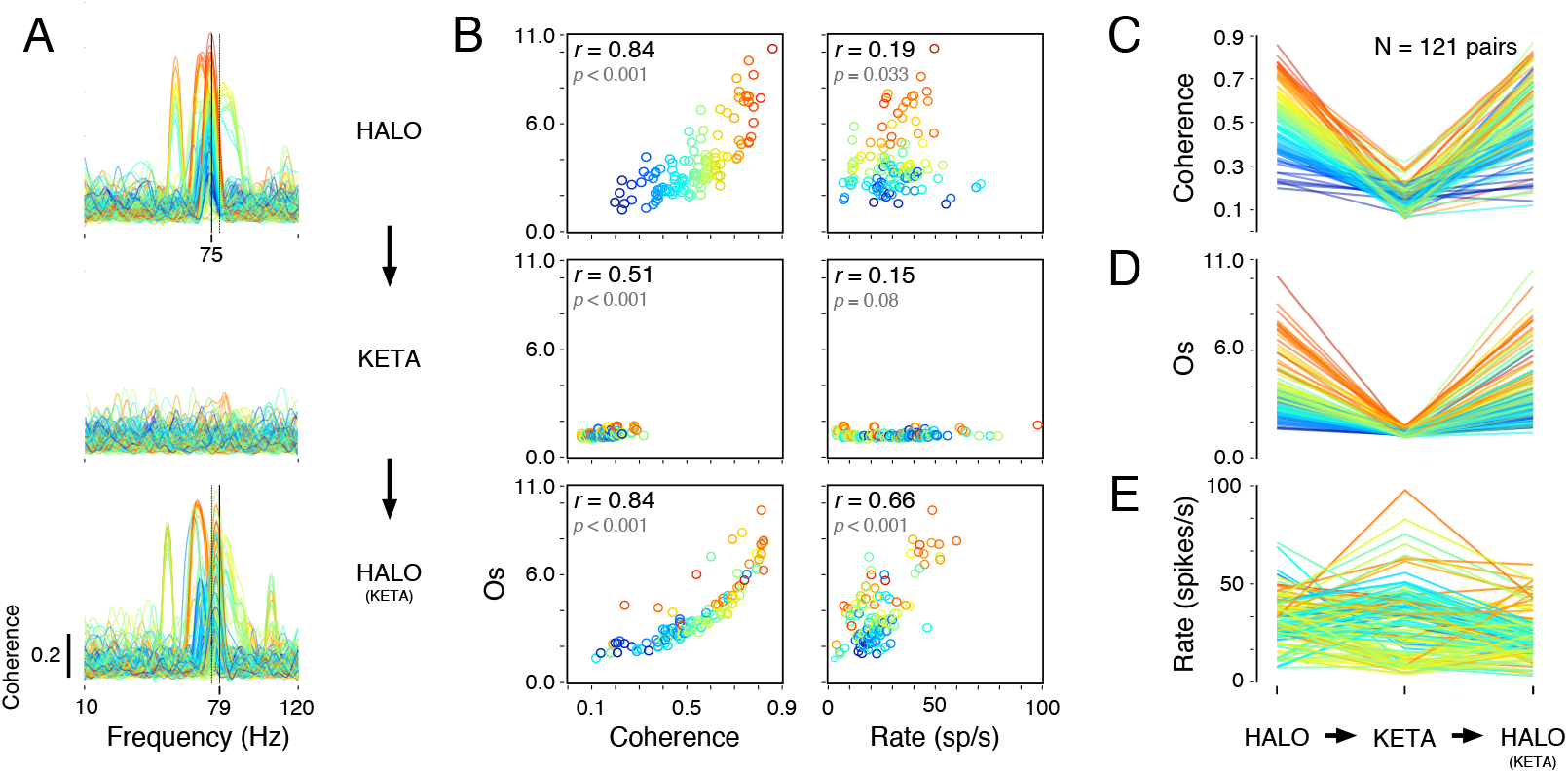
Halothane discontinuation produces a drastic reduction of gamma coherence. Data was collected from 121 pairs of single-cells in the retina (N= 13) and the LGN (N= 108 pairs) during a HALO → KETA → HALO (KETA) anesthesia transition protocol. Halothane, 1%. Same light stimulus was used across conditions. **(A)** Coherence functions computed for each cell pair. The functions are color-coded according to their peak amplitude measured in the HALO condition and plotted on a linear scale ranging from 0.1 (*purple*) to 0.9 (*red*). Mean frequency at coherence peaks are indicated by the continuous vertical lines (dotted lines serve a reference for comparisons). Note that after halothane anesthesia is reintroduced, gamma synchronization is restored. **(B)** Relationship between coherence and Os (*left column*) and rate and Os (*right column*). Pearson correlation (*r*) and its associated significance level (*p*) are indicated in the left-upper corner of each panel. Same color-code scheme as in (A). **(C)** Line plot of coherence changes over the HALO → KETA → HALO (KETA) transition. **(D)** Same as in (C), but for Os. **(E)** Same as in (C), but for firing rates. Unlike coherence and Os, there is no systematic changes in firing rates when halothane is removed.

Figure 5C-D shows the effect of halothane removal on the coherence and OS metrics for each cell pair. This effect was highly significant (coherence, T2 = 105.8, *p* < 0.0001; Os, T2 = 182.1, *p* < 0.0001; Friedman test). The coherence median was 0.54 (IQR = 0.22) for HALO, got reduced to 0.15 (0.08) for KETA, and returned to 0.54 (0.26) for HALO (KETA). This reduction amounts to a factor of 3.6. The difference in coherence across both HALO # KETA and KETA # HALO (KETA) transitions were significant (*p* < 0.0001, *p* < 0.0001, respectively; Wilcoxon signed-rank test with Bonferroni correction). Similar highly significant differences were also verified for Os. The Os median was 3.39 (IQR = 2.10) for HALO, got reduced to 1.24 (0.16) for KETA, and returned to 3.36 (2.14) for HALO (KETA). Firing rate modulation, on the contrary, showed high variability but no systematic decrease when halothane was removed (Figures 5E, median for the HALO and KETA groups = 31.5 and 32.0 spikes/s, respectively; *p* = 0.63, Wilcoxon signed-rank test). For a few pairs, there was even a sharp increase in responsiveness. Thus, a reduction in firing rate is unlikely to explain the drastic waning of gamma responses in the KETA condition.

By the end of the HALO # KETA # HALO (KETA) protocol, most cell pairs regained coherence values similar to those that appeared initially. There were no significant differences in coherence between HALO and HALO (KETA) conditions (*p* = 0.25, Wilcoxon signed-rank test). Only one from 121 cell pairs did not return to its original coherence confidence level when halothane was restored. Analysis of frequency measured at the coherence peak reveals a similar behavior (Supplemental Figure 1A-C). Moreover, the mean phase angle and phase-locking estimate across trials (PLV) also tended to be maintained after halothane restoration (Supplemental Figure 1D-F). A *post-hoc* Wilcoxon signed-rank test ran between the population medians of coherence phase angle for HALO (median = 11.13°, IQR = 52.79°) and for HALO (KETA) (9.56°, 43.85°) showed no significant difference (*p*= 0.1407). Interestingly, the phase relationships for each cell pair was strikingly consistent across trials for HALO (median = 0.88, IQR = 0.155) and resumed a similar profile for HALO (KETA) (0.88; 0.165). This similarity was confirmed statistically (*p* = 0.089).

In control experiments, we checked whether adding ketamine could interfere with the gamma oscillations induced by 1% halothane (Supplemental Figure 2). For this transition, instead of replacing halothane with ketamine, we added ketamine (10 mg/ Kg) to halothane (HALO + KETA) in cats that had been maintained under halothane for many hours (HALO-Only). From this analysis, it is clear that ketamine has no influence on the gamma responses induced by halothane.

### No evidence of retinal gamma in awake cats

If the pharmacological action of halothane is the primary cause of gamma oscillations in the retina, these oscillations should not be observed in awake animals. To verify this point, we set out to record from the LGN of awake cats, free of anesthetics. During such experimental sessions, the cats were subsequently anesthetized by ketamine and then by halothane (AWAKE # KETA (ONLY) # HALO (KETA) protocol). The recording electrodes were kept in place so that neuronal responses of the same groups of cells (MUA) could be directly compared across states.

Figure 6 shows spike-field coherence data obtained for the 31 pairs of recording sites from 2 cats for which we were able to characterize neural responses across state transitions. The maintenance of isolated units across channels throughout this transition protocol was not always possible, particularly after halothane delivery. This is why we decided to use spike-field coherence for it requires only one isolated cell per pair of sites. For this dataset, we used the same large light disk stimulus as before. Average coherence functions (Figure 6A, upper panel) revealed a striking difference between AWAKE and KETA (ONLY) states, on the one hand, and HALO (KETA) on the other. This difference was even more prominent after alignment at peak frequency before averaging (Figure 6A, bottom panel). The absence of gamma responses in the awake state and under ketamine is evidenced by the fact that the distributions of coherence and Os estimates were close to zero (median = 0.11 and 0.10, respectively) and uncorrelated (*r* = −0.19 and 0.02 for awake and ketamine, respectively; Figure 6B). As expected, introduction of halothane led to a significant increase in both coherence and Os (*p* < 0.001, Wilcoxon Signed Rank test; Figure 6C-E). As shown before, gamma coherence changes were not a direct consequence of rate modulation.

**Figure 6.**
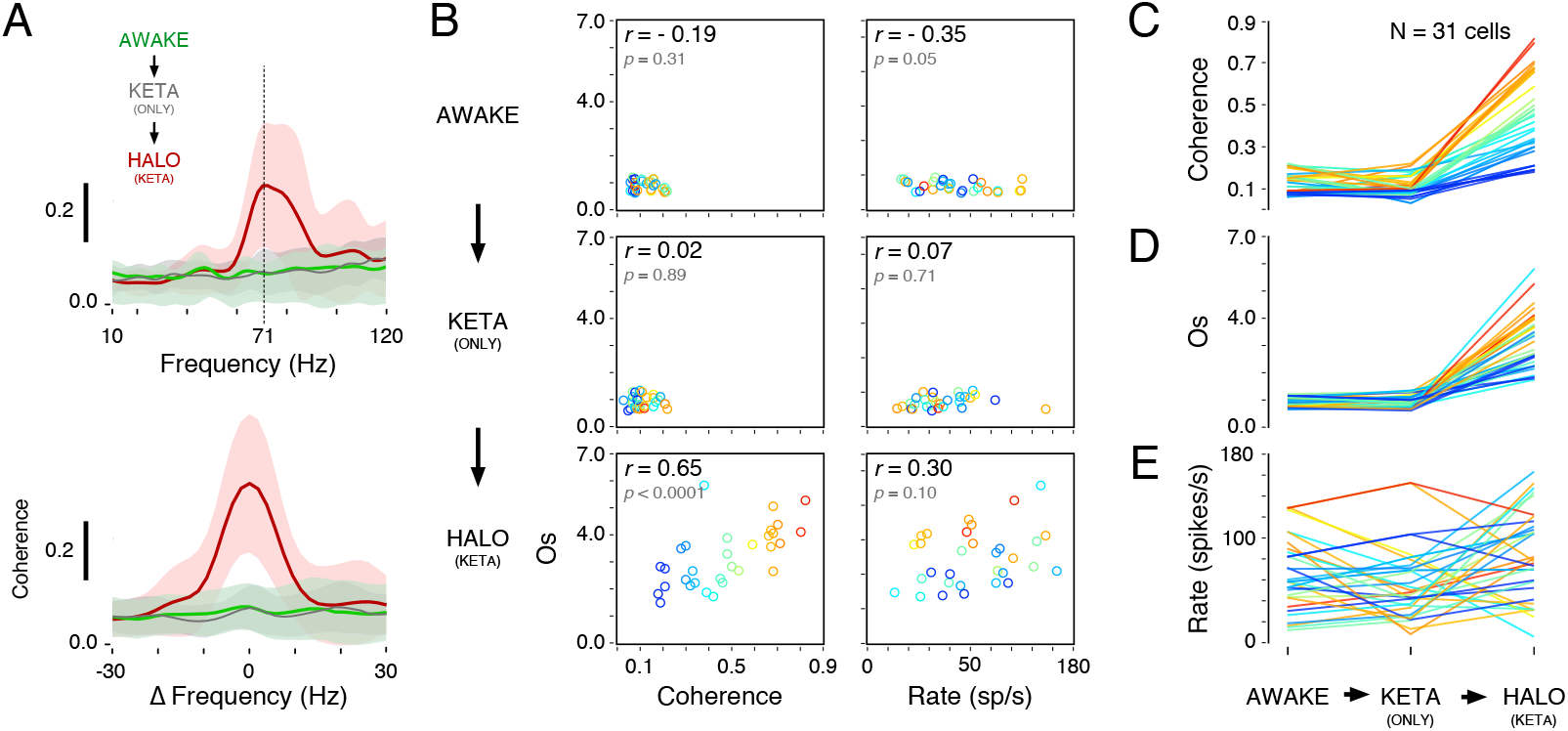
Gamma is absent in the awake state and appears only after halothane delivery. Recordings were performed in the LGN of awake cats, which were subsequently anesthetized by ketamine and then halothane. By keeping electrodes in place, responses of the same cell group could be compared across conditions (AWAKE → KETA (ONLY) → HALO (KETA) protocol. Data was obtained for 31 single cells. Stimulus, light disk displayed on a computer screen. **(A)** *Upper panel:* Average spike-field coherence function for the AWAKE (*green traces*), KETA (ONLY) (*gray*) and HALO (KETA) (*red*) conditions. Standard deviations of the mean are shown in lighter shadings. *Lower panel:* Same average spike-field coherence after being aligned by their respective peak frequency. **(B)** Relationship between coherence and Os (*left column*) and rate and Os (*right column*). Points are color-coded according to their peak amplitude measured in the HALO (KETA) condition ranging from 0.1 (*purple*) to 0.9 (*red*), as indicated in (C). **(C)** Line plot of coherence changes through the AWAKE → KETA (ONLY) → HALO (KETA) transition. **(D)** Same as in (C), but for Os. **(E)** Same as in (C), but for firing rates.

### Response entrainment

We often noticed a strong and stationary oscillatory component in the neuronal responses to large stimuli when cats were awake or under ketamine anesthesia (Figure 7). This component was entrained by the monitor vertical retrace (usually set at 100 Hz), as it disappeared after subtracting the shift predictor from the raw correlogram (Figure 7A). The main frequency component of the responses matched that of the monitor refresh, and remained unchanged during the whole course of the response. In contrast, under halothane, oscillations typically exhibited a pronounced decay in frequency after stimulus onset. In the example shown in Figure 7, oscillation frequency stabilized at ~80 Hz after an exponential decline starting at 93 Hz. When the shift predictor is subtracted, the gamma oscillations persist. Observe that the gamma peak is broader and has a lower amplitude in the HALO (KETA) as compared to the AWAKE and KETA (ONLY) conditions (Figure 7C). This is due to the significant decay in gamma frequency within the analysis window (500 ms).

**Figure 7.**
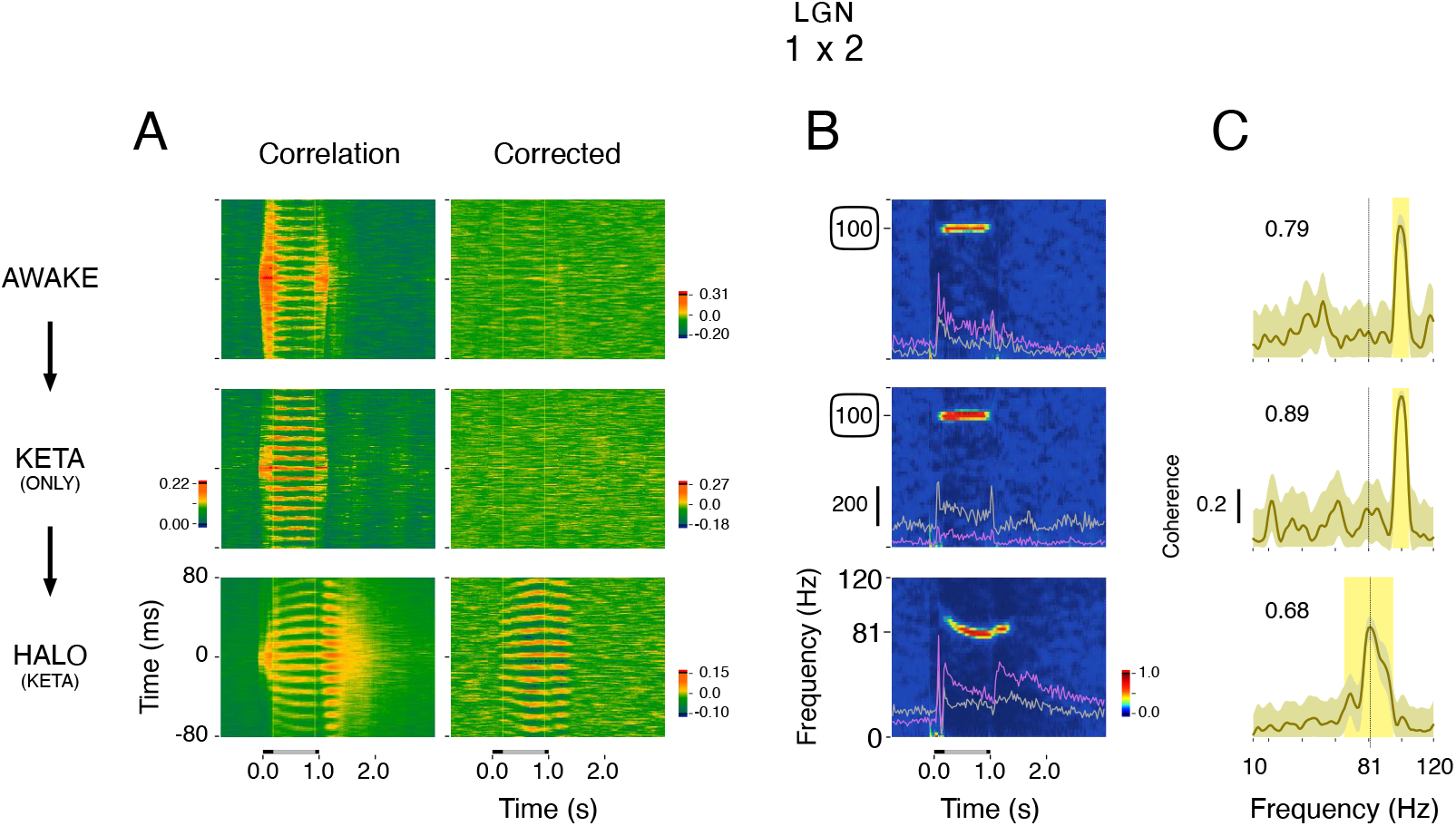
Example of response entrainment to the monitor refresh in the awake cat. MUA recordings were obtained from two separate electrodes placed in the LGN (lamina A) through the AWAKE → KETA (ONLY) → HALO (KETA) transition. For all three conditions, stimuli were presented on a computer monitor running at 100 Hz. **(A)** *Leftward column*: Sliding window cross-correlograms show strong response entrainment to the monitor refresh rate in the AWAKE and in the KETA (ONLY) condition. Notice that the entrainment frequency is constant over the whole presentation of the light stimulus. For the HALO condition, responses show strong gamma oscillation with a characteristic decay of frequency. Observe that the responses to offset of the light stimulus are also oscillatory at a slightly higher frequency. *Leftward column*: Sliding window cross-correlograms after shift-predictor correction. **(B)** Cross-correlation time-frequency plots. PSTH scale bar, 200 spikes/s. Gamma peak frequency was 81 Hz for the (HALO (KETA) condition. The round boxes indicate the monitor refresh rate (in this example, 100 Hz). **(C)** Coherence functions. Response entrainment results in a high amplitude, narrow coherence peak (0.79 and 0.89 for the AWAKE and in the KETA (ONLY) conditions, respectively). In the HALO (KETA) condition, gamma peak was relatively broader and had a lower amplitude (0.68).

As a control for response entrainment, we used LEDs to generate flicker-free, uniform light patch stimuli. Typically, in awake cats, stimulation by LED did not evoke rhythmic patterning in the responses, as opposed to CRT monitor stimulation, which often did so (Figure 8A). The same phenomenon was seen under ketamine (not shown). The spike-field coherence data shown in Figure 8B was consistent for all pairs of sites in the same recording session for which we were able to run the LED *vs*. CRT protocols. In awake cats, spike-field coherence was significantly higher for monitor displayed stimuli than for LED (*p* < 0.0001, Wilcoxon rank sum test); under halothane, no statistical difference was detected (*p* = 0.83). This control experiment rules out the possibility that rhythmic responses are generated internally in the wake state (or under ketamine). Instead, they are caused by the external flickering of the stimulus.

**Figure 8.**
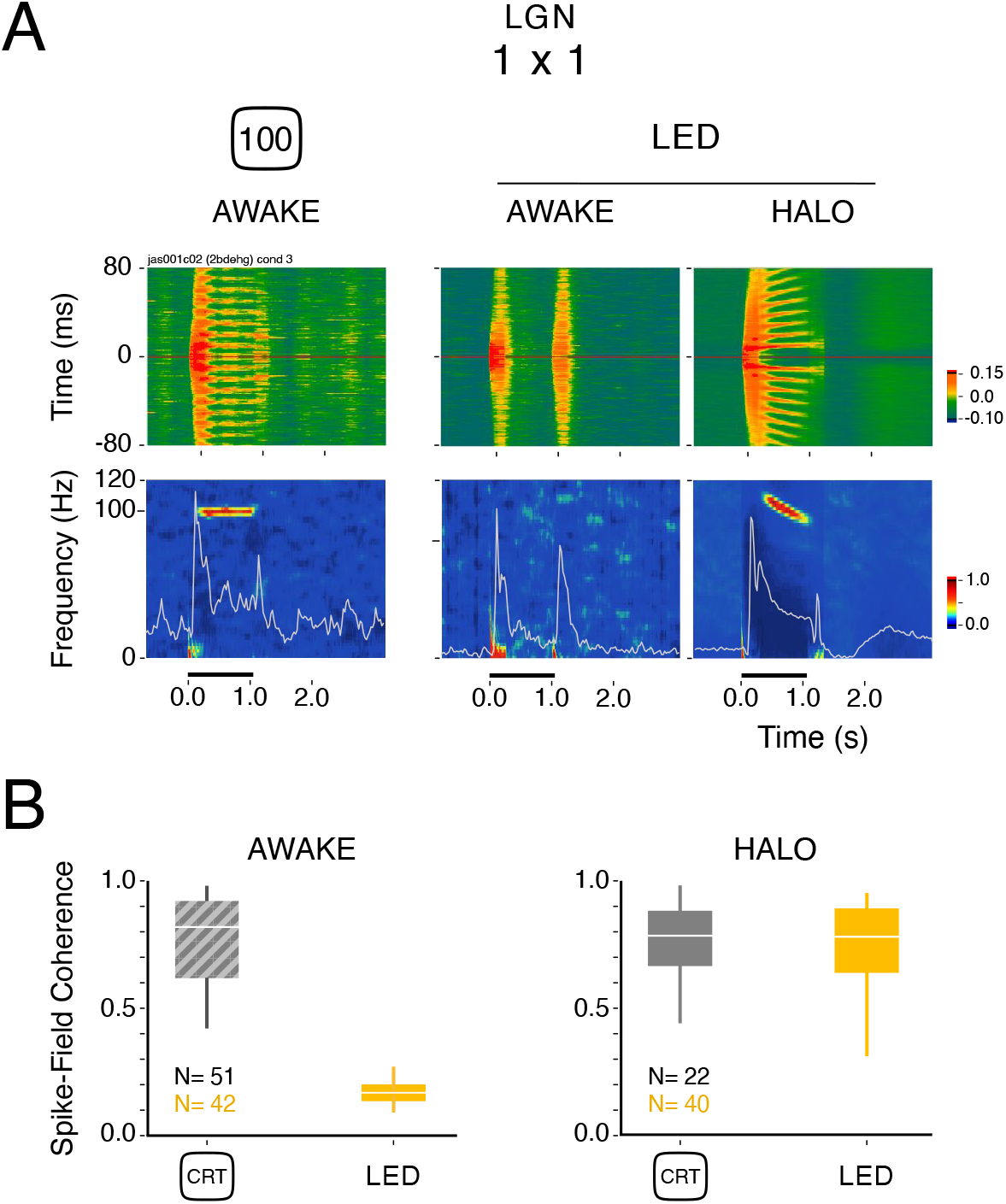
Comparisons of responses to CRT display and LED. **(A)** Autocorrelograms (*upper row)* and corresponding time-frequency plots (*bottom row*) computed for a group of cells (MUA) recorded in the LGN. In the AWAKE condition, the CRT monitor entrained the neuronal responses (100 Hz), while stimulation by LED generated no oscillatory components. In the HALO condition, however, the LED stimulus could induce strong gamma with characteristic frequency decay. **(B)** Population analysis of spike-field coherence median differences between CRT and LED stimulation in the AWAKE (N= 93 cells) and the HALO (N= 62) conditions.

Notably, under halothane, neither the frequency nor the strength of gamma coherence were affected by the flickering of the CRT monitor (Figure 9). We characterized the responses to a large light disk as a function of five CRT refresh rates (60, 75, 85, 100 and 120 Hz). Data from two recordings of the same cat were analyzed for awake and anesthesia states (cells were included only if exhibiting response entrainment in the awake state). Oscillation frequency was obtained for two successive analysis windows (Win1, Win2). In awake states (Figure 9A), the cells always followed the refresh of the screen. Mean peak frequency remained constant throughout a trial (Win 1 *vs*. Win2, *p* = 0.21, Wilcoxon matched-pairs signed rank test), indicating that the oscillations observed in the awake state were completely determined by the screen refresh. Under halothane anesthesia (Figure 9B), on the other hand, monitor flicker did not influence the characteristic frequency-decay profile of gamma oscillation. The oscillation frequency values estimated during the early part of the responses (Win1) were significantly higher than those during the late epoch (Win2) of the responses (p < 0.0001, Wilcoxon matched-pairs signed rank test). Irrespective of the screen refresh rate, the distribution of oscillation peak frequencies remained clustered around a median of 84.0 Hz (Win1, IQR = 9.7) and 76.2 Hz (Win2, IQR = 3.9). As shown in the examples, both entrainment and gamma coherence amplitudes were comparable, even at the highest monitor refresh (120 Hz). Most importantly, in our data, we did not find a single example of co-occurrence of entrainment of the responses by the monitor refresh and internally generated gamma (under halothane). This result suggests that, within the gamma band, the two rhythmic response modes cannot coexist in the retina, as proposed by Rager and Singer (1998) in the visual cortex (but see Duecker, 2021).

**Figure 9.**
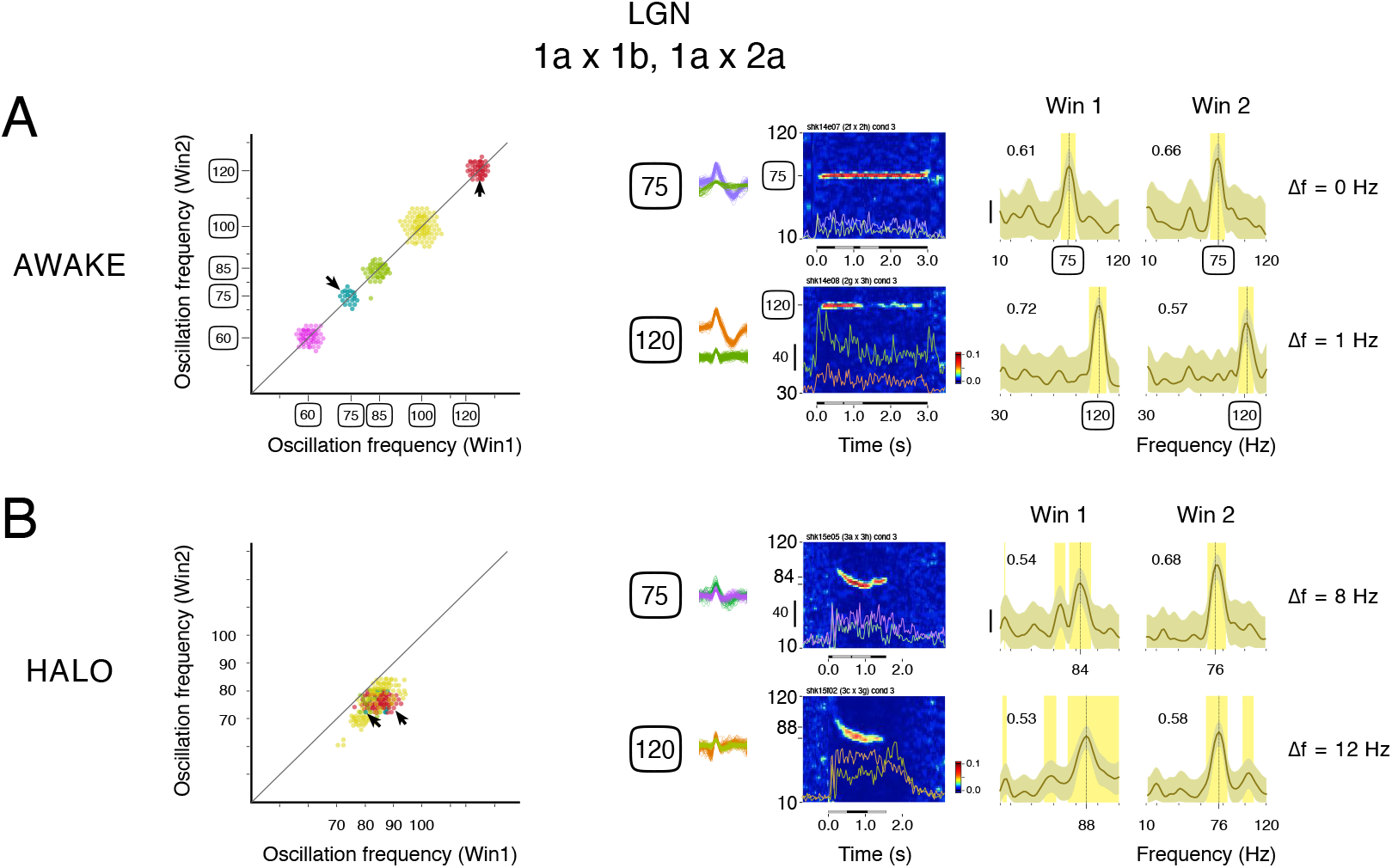
Gamma synchronous oscillations induced by halothane supersede stimulus entrainment. Population analysis of LGN single-cell responses to a light stimulus displayed on a CRT monitor with various refresh rates (60, 75, 85, 100, 120 Hz). **(A)** AWAKE condition (N= 265). Scatter plot of oscillation frequency computed for two subsequent analysis windows (Win1 and Win2, 500 ms). *Right panels:* examples of time-frequency plots and coherence functions corresponding to the data points identified by the arrowheads on the scatter plots. Coherence functions are shown for Win1 and Win2, as indicated on the time-frequency plots. In the examples, monitor refresh rate was set to 75 and 120 Hz. **(B)** HALO condition (N= 209), same conventions as in (A).

### Grand comparisons

In Figure 10, we show the relationship between gamma coherence and Os for all the single-cell data obtained in our study (5,392 cell pairs from a same electrode; 7,244 across electrodes; 9 cats). Comparisons were made between AWAKE and halothane conditions (HALO and HALO (ONLY), see Experimental Design in Material & Methods) as function of stimulation method (CRT monitor or LED). Cell pairs that were entrained by the monitor were sorted from those that showed no entrainment. In the AWAKE state, high values of coherence and Os were seen only when the responses were entrained externally by the monitor. For the cell pairs showing no entrainment, gamma coherence and Os values were consistently low. Accordingly, the controls made with LED stimuli showed no signs of gamma for the awake state. Under HALO, on the other hand, we observed the same positive relationship between coherence and Os as seen before for the anesthesia transitions (see Figure 5). This behavior may be considered as a hallmark of gamma synchronization in the retina, suggesting that oscillation in the gamma range is a necessary condition for synchronization.

**Figure 10.**
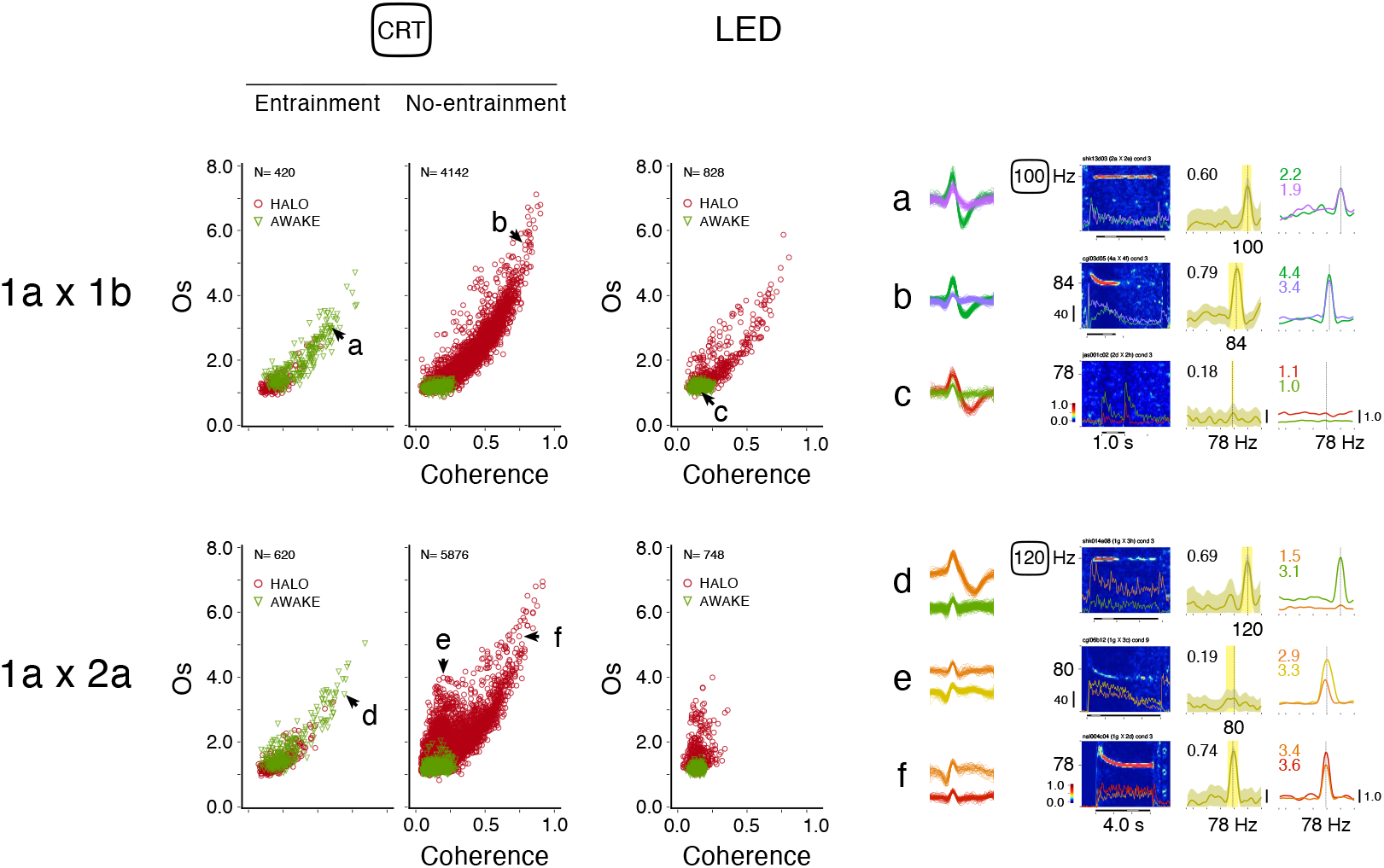
Grand comparisons of gamma coherence and oscillation strength for the awake and the halothane conditions. Relationship between coherence and Os for all pair combinations derived from the entire sample of single-cells recorded in the retina (N= 180) or in the LGN (N= 2261) obtained in 9 cats. Data was split by states (AWAKE and HALO) and stimulation type (CRT and LED). Cases of response entrainment are presented separately. *Upper panels:* same-electrode pairs (N= 5392). *Lower panels:* cross-electrode pairs (N= 7244). *Right panels:* examples of time-frequency plots, coherence functions and power spectra for individual channels corresponding to the data points identified by the letters (**a** to **f**) on the scatter plots. Os values are shown in the power spectrum plots. A shown in example **e**, an oscillatory profile in both power spectra does not necessarily result in a high coherence. Most likely, these cases represent cell pairs that receive inputs from different eyes.

Note, however, that this relationship showed deviant points. This occurred mostly for cross-electrode pairs. As one can see in the example in Figure 10 (point **e**), low coherence values were found even when the power spectra computed for each channel individually had an oscillatory profile. A plausible explanation for this paradoxical finding is that these cases correspond to pairs of cells receiving inputs from different eyes. As shown by Neuenschwander and Singer (1996) correlated activity in the LGN was strong only for cell pairs with inputs from the same eye. Importantly, in our study, entrained responses had the same overall characteristics as the gamma generated internally under halothane: narrow peaks and strong correlation between coherence and Os. These trends were seen both for pairs of single-cells from the same electrode and across electrodes. Overall, these findings suggest that stimulus entrainment and gamma oscillations in the retina may share the same synchronization mechanism.

Figure 11 presents single-cell coherence data sorted by significance (95% confidence interval) using the same grouping as Figure 10. The proportion of significant values varied across groups (likelihood ratio χ^2^ test, χ^2^_5_ = *p* < 0.0001; pairs from the same electrode) and was positively correlated with within-group medians (Pearson *r* = 0.94, p < 0.0057). Only 45.5% of cell pairs (167 of 367; median = 0.14, IQR = 0.06) in the non-entrain group had significant values for the AWAKE state, compared to 87.6% (3307 of 3775; median = 0.39, IQR = 0.24) for HALO. An inverse trend was seen for the entrainment group, as 76.6% of cell pairs (203 of 265; median = 0.40, IQR = 0.23) had significant values for the AWAKE state, compared to 49.0% (76 of 155; median = 0.22, IQR = 0.08) for HALO. These differences in proportion were well above the Marascuillo’ critical range value of equality (non-entrainment: absolute proportion difference = 0.42, critical range = 0.074; entrainment: absolute proportion difference = 0.27, critical range = 0.16). Within both the *No-entrainment* and the *Entrainment* groups, differences in coherence medians between AWAKE and HALO conditions were equally highly significant (*p* < 0.0001, Wilcoxon-rank test). Notably, when comparing the distributions between *No-entrainment* + HALO and *Entrainment +* AWAKE groups, there is no significant difference in coherence (*p* = 0.1142, Wilcoxon-rank test). In the LED control group, coherence values were similarly distributed to those in the non-entrainment group. Despite similar trends, the differences between cross-electrode pairs are smaller than those observed between same-electrode pairs. Overall, these results reinforce that, in the awake state, relatively high coherence levels always reflect entrainment of the responses to external rhythmic inputs.

**Figure 11.**
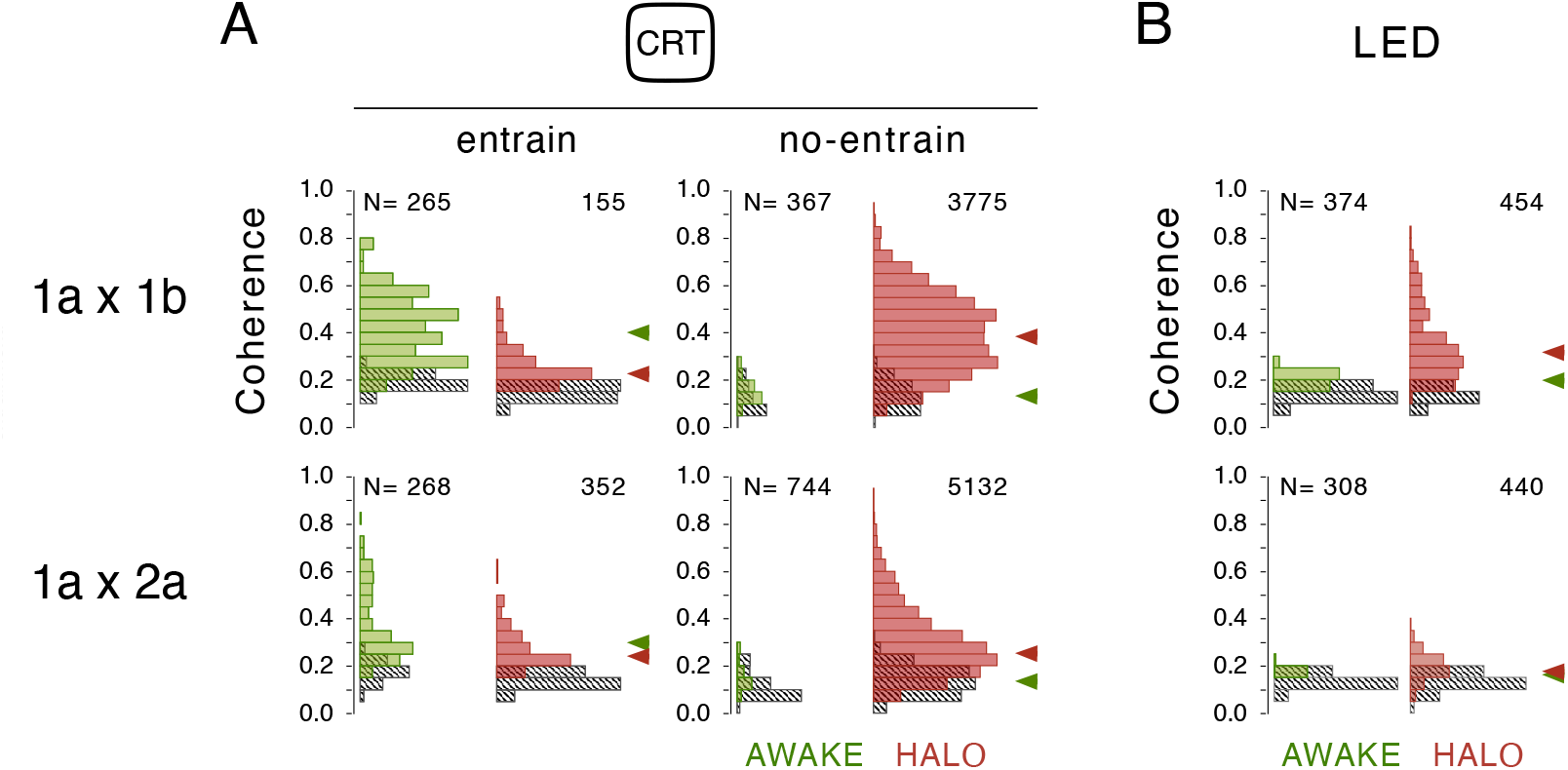
Summary statistics of coherence for gamma oscillations *versus* response entrainment. Same data set as in Figure 10. Hashed bar histograms indicate cell pairs for which coherence was below the 95% significant level. Arrow heads indicate population medians. In the awake state, high coherence values are only found for cases of response entrainment. These values are comparable to those found for cell pairs under halothane with no entrained activity.

To better visualize the differences in gamma between awake and halothane states, we ranked the coherence functions obtained for all single cells by their frequency peak (4142 same-electrode cell pairs; and 5876 cross-electrode cell pairs). Pairs showing entrainment to the monitor were excluded. This analysis, presented in Figure 12, shows conclusively the absence of gamma in the awake state. In contrast, under halothane, gamma is always present in most of the data, exhibiting coherence functions with single, narrow peaks. Oscillation frequencies were consistent in the same-electrode and cross-electrode groups (median = 78.1 and 76.1 Hz, respectively).

**Figure 12.**
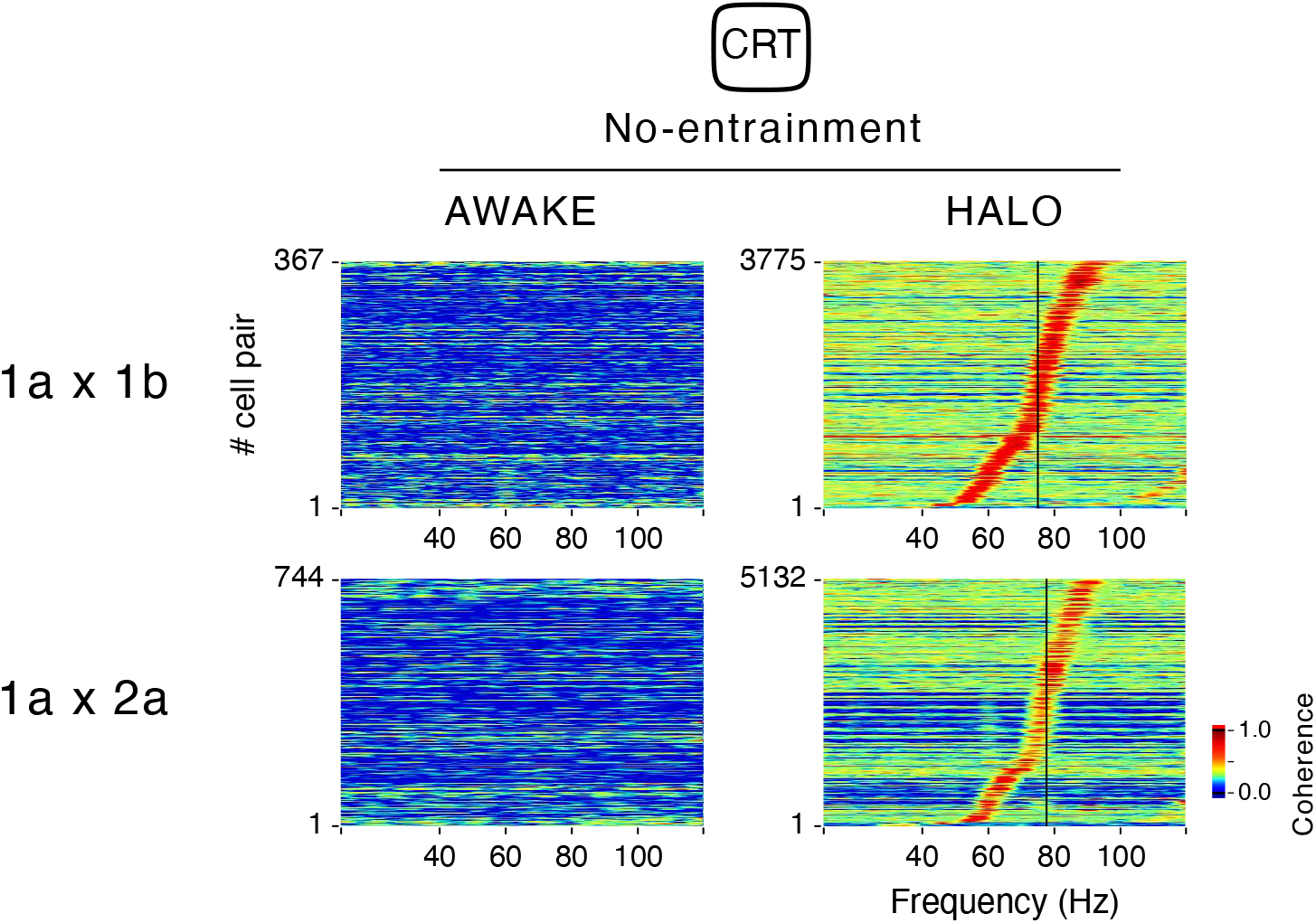
Compilation of coherence functions split between awake and halothane conditions for all single-cell pairs obtained in this study. Coherence functions obtained for all single-cell pairs (same-electrode, N= 4142; cross-electrode, N= 5876) are ranked by their frequency peak (taken from the cross-spectrum) and stacked one on top of the other as lines color-coded by amplitude (see *Z-scale*). Pairs showing response entrainment were excluded. Notice that, in the AWAKE condition, gamma is completely absent. Gamma is always present in most data under halothane, showing coherence functions with single, narrow peaks. Oscillation frequency median = 78.1 and 76.1 Hz for the same-electrode and cross-electrode groups, respectively.

In summary, in awake cats, the retinogeniculate system lacks robust intrinsic, visual gamma-band oscillations, but exhibits remarkable sensitivity to visual inputs flickering in the same frequency range.

## Discussion

Several lines of experimental (Ishikane et al., 2005; Ito et al., 2010; Koepsell et al., 2009; Neuenschwander and Singer, 1996; Roy et al., 2017) and theoretical evidence (Koepsell et al., 2010; Stephens et al., 2006; Warner and Sommer, 2020) support the hypothesis that gamma oscillations in the retina and LGN contribute to early visual processing. The goal of this study was to validate this hypothesis for natural vision. Under halothane (or isoflurane) anesthesia, gamma oscillations were robust and feature-specific, even for time-varying visual stimuli such as natural scene movies. However, to our surprise, gamma power in the retina was found to be inversely proportional to halothane concentration levels. This is the first time that such a relationship has been established. Moreover, we found no evidence of gamma activity in the LGN of awake cats, raising doubts about its functional meaning. In awake states, responses were often entrained to the monitor refresh (up to 120 Hz). We argue that response entrainment may be mediated by the same synchronizing mechanism that underlies the intrinsic retinal oscillations induced by halothane anesthesia.

GABAergic interneurons are known to play a critical role in the generation and control of gamma-band oscillations (Buzsáki and Wang, 2012; Cardin et al., 2009; Chen et al., 2017; Tiesinga and Sejnowski, 2009; Veit et al., 2017). In the vertebrate retina, many amacrine cells release GABA (Pourcho and Goebel, 1983; Yang, 2004), providing inhibitory inputs to bipolar cell terminals, retinal ganglion cells, and other amacrine cells (Diamond, 2017; Franke and Baden, 2017; Zhang and McCall, 2012; Zhang et al., 2004). Volatile anesthetics, such as halothane and isoflurane, among their various excitability control functions, primarily potentiate GABAA receptor activity (Franks, 2008; Sousa et al., 2000). Thus, halothane (or isoflurane) could facilitate inhibitory feedback loops at multiple levels in the inner retina, either pre- or postsynaptically. An increase in recurrent interactions within the highly interconnected inner plexiform layer (IPL) network could explain the massive synchronization of oscillatory activity exhibited by the ganglion cells. In line with this hypothesis, GABAA receptors seem to play an essential role in generating retinal oscillations. In the bullfrog, the GABAA antagonist bicuculline disrupts the oscillatory behavior of spiking discharges (Arai et al., 2004; Ishikane et al., 1999; Qiu et al., 2016). Oscillatory potentials recorded from the eye of the mudpuppy are also damped by bicuculline in a dose-dependent fashion (Fig. 3 in Wachtmeister, 1980).

Interestingly, the blockade of GABAA receptors suppresses the synchronization of oscillatory responses across long distances but preserves local, non-oscillatory interactions (Fig. 8 in Ishikane, 1999). Such interactions are abundant in the IPL (Arnett and Spraker, 1981; Brivanlou et al., 1998; DeVries, 1999; Mastronarde, 1989). Accordingly, it has been estimated, for example, that 15–30% of the spikes generated by individual ON-parasol cells consists of synchronous firing from large, spatially contiguous cell groups (Shlens et al., 2009). These interactions are mediated by gap junctions between amacrine and ganglion cells (Field and Chichilnisky, 2007; Hu and Bloomfield, 2003). Using a stimulus paradigm similar to Neuenschwander and Singer (1996), Roy et al. (2017) found that synchronous oscillations across long distances occur for responses to large, continuous stimuli, but not for disjointed spots restricted to the receptive fields. This phenomenon is mediated by wide-field polyaxonal amacrine cells. Moreover, the blocking of gap junctions causes suppression of long-range synchrony of oscillatory responses (Qiu et al., 2016; Roy et al., 2017), indicating that gap junction coupling in the IPL is a necessary condition to maintain the oscillations. Our experiments, in line with others (reviewed in Neuenschwander et al., 2002), provide strong evidence for long-range synchrony in the cat retina. Thus, junctional coupling is likely to be preserved under halothane (or isoflurane), although some evidence shows that GABAergic anesthetics may suppress gap junction function (Voss et al., 2010; Wentlandt et al., 2006).

At this point, it is worth emphasizing that all anesthetic agents used in studies reporting gamma oscillations in the retina or the LGN primarily act on GABAA receptors. Those include volatile anesthetics (Ariel et al., 1983; Arnett, 1975; Ito et al., 2010; Neuenschwander and Singer, 1996; Storchi et al., 2017), barbiturates (Bishop et al., 1964; Doty and Kimura, 1963; Laufer and Verzeano, 1967; Steinberg, 1966) and propofol (Koepsell et al., 2009). In our study, we used ketamine as non-gabaergic anesthetic control (Thomson et al., 1985). In contrast to halothane, no oscillatory responses were observed under ketamine. These findings were unexpected as volatile anesthetics diminish or completely suppress gamma rhythms in the visual cortex and the hippocampus (Faulkner et al., 1998; Xing et al., 2012) while ketamine increases gamma power (Li and Mashour, 2019; Mashour, 2014). Thus, gamma oscillations in the retina may depend on the recruitment of inhibitory synaptic inputs specifically mediated by GABAA-receptors.

Given that retinal gamma oscillations are absent in the awake cat, and presumably contingent upon GABAergic anesthesia, such oscillations should be considered as artifactual. This view is consistent with a recent study on cortico-thalamic interactions in the behaving monkey that found no evidence of gamma-band activity in the LGN (Bastos et al., 2014). Notwithstanding, this conclusion does not seem to apply to all vertebrate species. In isolated retina preparations, oscillatory responses have been reported for specific cell classes in frogs (dimming detectors; Ishikane et al., 1999) and mice (ON-alpha-cells; Roy et al., 2017). A narrowband oscillation in the retina and the LGN has also been identified in the behaving mouse, and linked with the activity of melanopsin photosensitive ganglion cells (Storchi et al., 2017). This cell type has a similar distribution in cats as in mice (Semo et al., 2005). However, it is unlikely that this system accounts for the gamma oscillations described here. The melanopsin-related narrowband oscillations in the mouse depend on high levels of illumination (Storchi et al., 2017), while the retinal gamma oscillations in the cat are maintained both at scotopic and photopic levels (Laufer and Verzeano, 1967; Neuenschwander et al., 1999).

In mice, the narrowband rhythm described in the LGN by Storchi et al. (2017) is similar to that of Neill and Stryker (2010) in V1. Using optogenetics, Saleem et al. (2017) demonstrated that this V1 rhythm actually originates in the LGN, which led them to propose an homology with the gamma oscillations in the anesthetized cat. We consider this hypothesis unlikely because the rhythm reported in the mouse features a much narrower bandwidth (~ 60 Hz) when compared to the retinal oscillations in the cat (40 to 120 Hz). Besides, the frequency reported in the mouse is stationary over time, which contrasts with the exponential decay described in our study (see examples in Figure 3). Interestingly, Storchi et al. (2017) also found narrowband oscillations under isoflurane. Like in our data, a frequency decay is evident under isoflurane anesthesia but not in the awake condition (Storchi et al., 2017, Fig. S1H and J, respectively). Thus, the narrowband 60 Hz oscillation in the behaving mouse may represent a rhythm entirely distinct from that induced by anesthesia.

In the awake cat, neuronal responses were often entrained to the monitor refresh for test frequencies as high as 120 Hz. Response entrainment depended on visual drive and was not seen in spontaneous activity. It was most robust for large, high-contrast stimuli, consistent with previous studies in the retina, LGN, and cortex of anesthetized cats (Eysel and Burandt, 1984; Neuenschwander et al., 1999; Wollman and Palmer, 1995). Phase-locking of spiking responses to flickering stimuli has been found in several species other than cats, including anesthetized three shrews and monkeys (Veit et al., 2011; Williams et al., 2004), as well as alert monkeys (McClurkin et al., 1991) and humans (Duecker et al., 2021; Krolak-Salmon et al., 2003). Notably, in our study, the entrained responses were generally superseded by the internally generated oscillations, even when the monitor refresh was several tens of Hz higher or lower than the intrinsic frequency (Figure 9). It seems that retinal oscillations induced by anesthesia behave like an attractor insensitive to the temporal structure of the stimulus.

Several aspects of response entrainment are similar to intrinsic gamma oscillations. First, both rhythms are dependent on stimulus properties such as size and contrast (Neuenschwander et al., 1999; Williams et al., 2004). Second, peak frequencies for intrinsic (mean 80 ± SD Hz; max. 118 Hz) and entrained rhythms (tested up to 120 Hz) can reach relatively high values, well above the temporal resolution of both the X- and Y-cells in the cat LGN (12 and 20 cycles/s, respectively; (Lehmkuhle et al., 1980). Equally important, retinal oscillations are strongly phase-locked to stimulus onset for the first few hundred milliseconds, even for a non-flickering stimulus (Neuenschwander et al., 1999). This response behavior has often been overlooked, even though stimulus-locking is robust and can be easily detected when response histograms are plotted at high resolution (Fig. 5 in Neuenschwander et al., 1999; Fig. 2 in Qiu et al., 2016; Fig. 2 B in Storchi et al., 2017; Fig. 5E in Roy et al., 2017). It can also be easily seen in electroretinogram traces (Storchi et al., 2017; Wachtmeister, 1980). Eventually, these highly phase-locked signals reach the cortex. This feedforward signal propagation could explain the paradoxical findings of Imas et al. (2004). In their study, gamma-evoked potentials in the cortex of the rat show an increase in power as a function of halothane levels. We believe that these evoked oscillations have an origin in the retina and not in the cortex.

Based on the above considerations, we propose that, like the retinal gamma oscillations, response entrainment requires active synchronization, and do not merely reflect a passive phase-locking of individual cell responses. As discussed above, this synchronization mechanism is probably mediated by gap junctions of wide-field amacrine cells at the IPL level, though coupling within the outer plexiform layer cannot be excluded (Bloomfield et al., 1995; DeVries et al., 2002; Greb et al., 2017; Vaney, 1994). Thus, just as a large light spot is capable of triggering long-distance synchronous oscillations, the spatiotemporal coherent flicker of a CRT screen also evoke active, mass interactions of cell responses. If true, several predictions can be made. First, small, local neuronal populations in the retina, when activated in isolation, should not be able to follow fast flickering stimuli, just as small spots do not trigger synchronous oscillatory responses (Roy et al., 2017). Second, blockage or deletion of gap junctions should interfere with the response entrainment. Third, bipartite, concentric stimuli flickering over receptive fields in a center-surround incongruous manner should not result in response entrainment.

From the viewpoint of a readout mechanism, there should be no difference between an internally generated oscillation in the retina and an externally entrained rhythm. We argue that visual processing is robust against the influence of such fast, steady-state oscillatory inputs. That is probably why the periodic flickering of visual displays does not interfere with image perception, even though the visual cortex is massively entrained by rhythmic inputs (Duecker et al., 2021; Lyskov et al., 1998; Rager and Singer, 1998; Williams et al., 2004). We emphasize that this conjecture does not conflict with neural code models based on non-oscillatory synchronization mechanisms in the retina (Baccus, 2007; Pillow et al., 2008; Shlens et al., 2008).

## Figure Legends

**Supplemental Figure 1.**
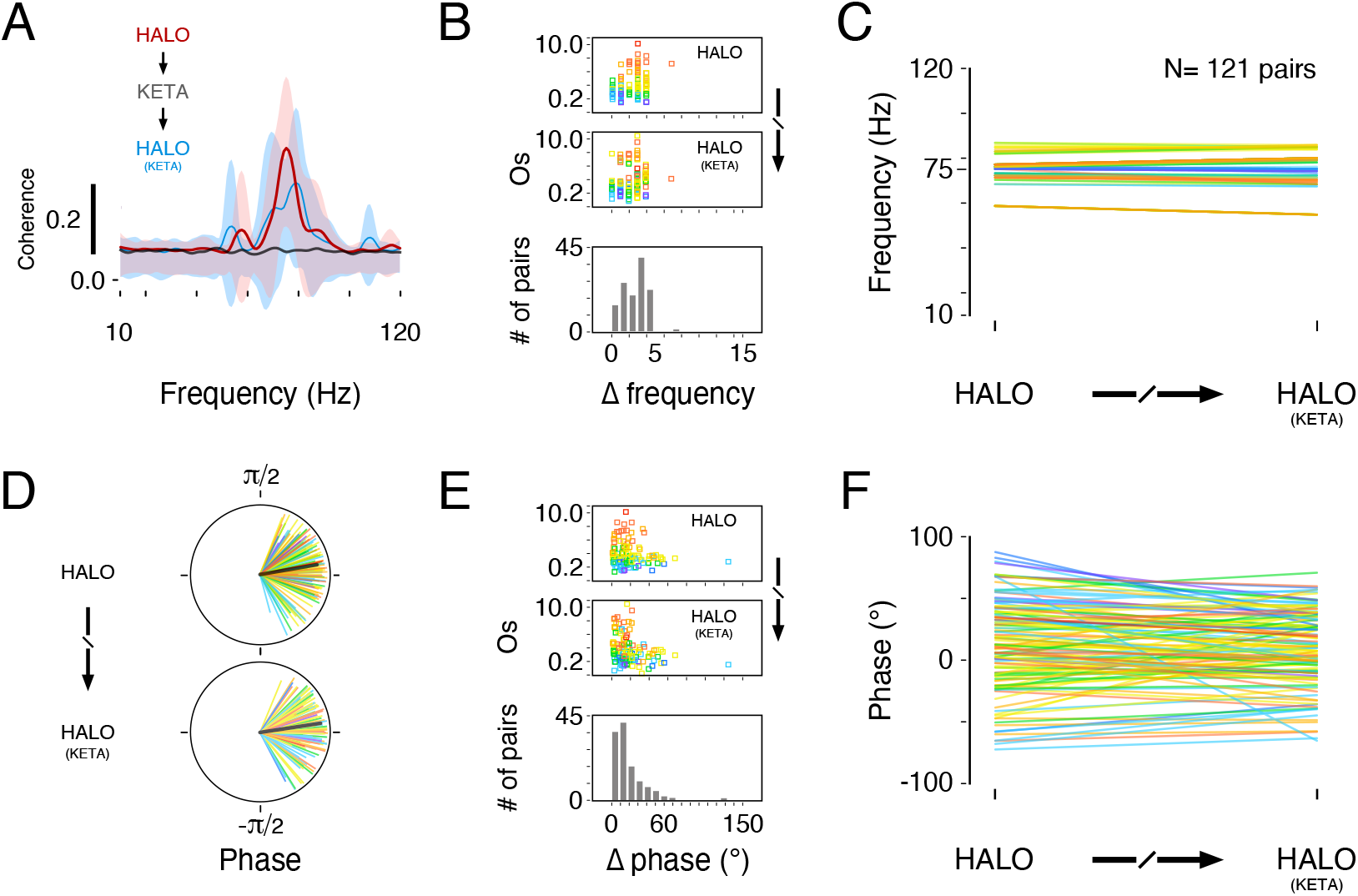
The transition protocol HALO → KETA → HALO (KETA) does not substantially change the frequency and phase characteristics of gamma oscillations after recovery. Same data as in Figure 5. **(A)** Average coherence functions for HALO (*red*), KETA (*gray*) and HALO (KETA) (*blue*) conditions. Standard deviations of the mean are shown in lighter shadings. **(B)** *Upper panels:* Scatter plots of Os as function of oscillation frequency differences between baseline (HALO) and recovery (HALO (KETA)) conditions. Points are color-coded according to the corresponding coherence value measured in the HALO condition (0.1, *purple;* 0.9, *red*). *Lower panel:* Counts of cell pairs according to frequency differences data plotted in the upper panels. In most data, differences are less than 5 Hz. **(C)** Line plot of frequency changes between baseline and recovery conditions. **(D)** Circular representation of mean phases computed from coherence analysis for each cell pair in baseline and recovery conditions. *Black line*, vector average across cell pairs. **(E)** Same as in (B), but for phase differences. **(F)** Same as in (C), but for phase values.

**Supplemental Figure 2.**
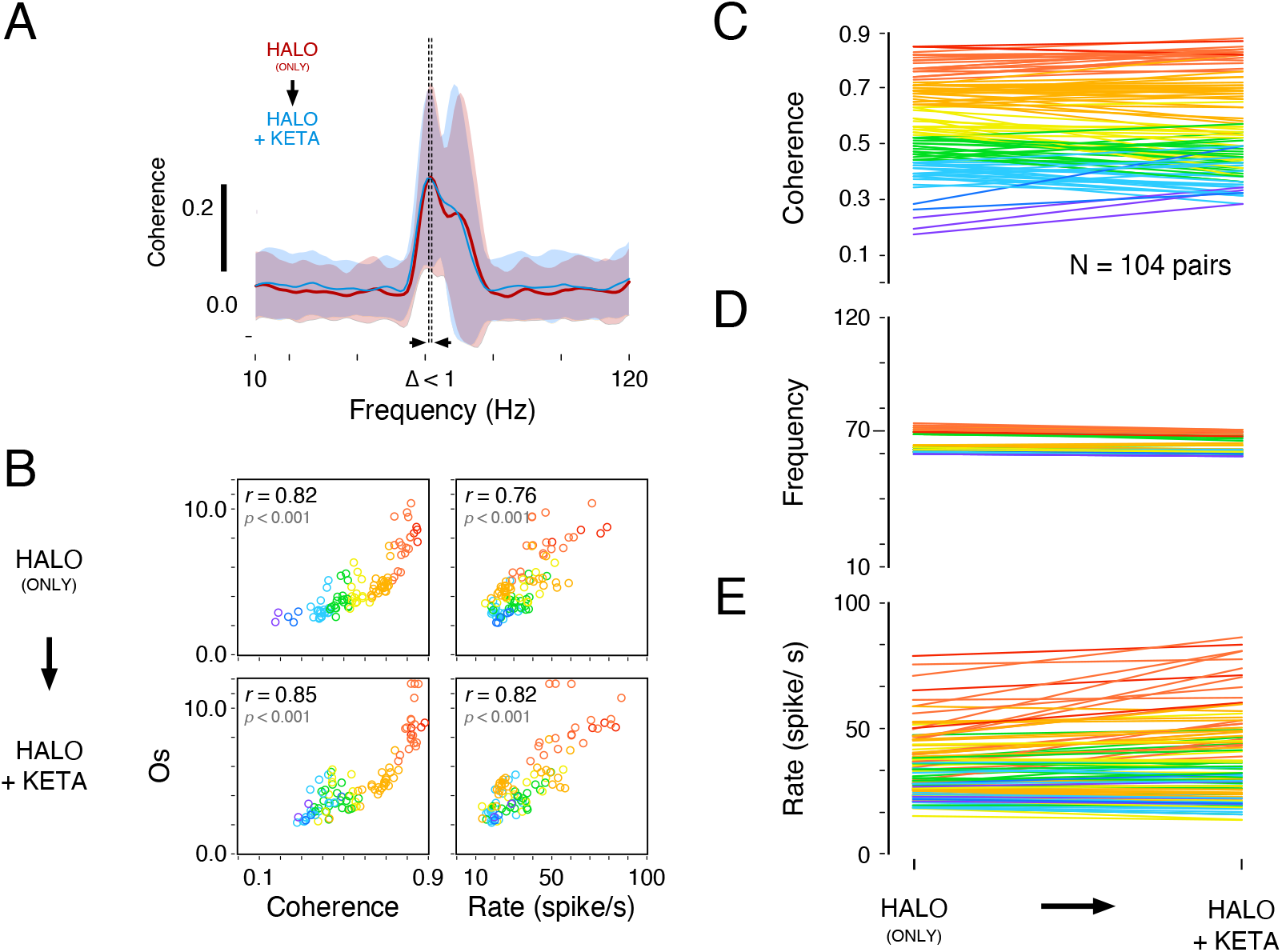
Ketamine does not interfere with halothane-induced gamma oscillations. In this protocol (N= 104 cell pairs), we added ketamine (10 mg/ Kg) to the anesthesia (HALO + KETA) in cats that had been maintained under halothane for hours (HALO (ONLY)). **(A)** Average coherence functions for HALO (*red*), and HALO + KETA (*blue*) conditions. Standard deviations of the mean are shown in lighter shadings. **(B)** Relationship between coherence *vs*. Os (*left column*) and rate *vs*. Os (*right column*). Pearson correlation (*r*) and its associated significance level (*p*) are indicated in the left-upper corner of each panel. Points are color-coded according to the corresponding coherence value measured in the HALO (ONLY) condition (0.1, *purple;* 0.9, *red*). **(C)** Coherence values before and after ketamine administration. Difference in coherence was not significant (*p* = 0.124, Wilcoxon sign-rank test). Medians, 0.56 and 0.53 for HALO (ONLY) and HALO + KETA, respectively. **(D)** Same as (C), but for coherence peak frequency. Difference in frequency was extremely small (1 Hz), though statistically significant (*p* < 0.0001, Wilcoxon sign-rank test). Medians, 62.5 and 61.5 Hz. **(E)** Same as in (C), but for firing rates. There was a much higher variability in rates as compared to the coherence and frequency values. Difference in rate median was statistically significant (*p* = 0.0014, Wilcoxon sign-rank test), with a slight increase (median 29.6 to 31.9 spikes/s, 8% increase) in rate after ketamine.

